# The neurophysiology of closed-loop auditory stimulation in sleep: a magnetoencephalography study

**DOI:** 10.1101/2022.12.19.521103

**Authors:** Hugo R. Jourde, Raphaëlle Merlo, Mary Brooks, Meredith Rowe, Emily B.J. Coffey

## Abstract

Closed-loop auditory stimulation (CLAS) is a brain modulation technique in which sounds are timed to enhance or disrupt endogenous neurophysiological events. CLAS of slow oscillation up-states in sleep is becoming a popular tool to study and enhance sleep’s functions, as it increases slow oscillations, evokes sleep spindles, and enhances memory consolidation of certain tasks. However, few studies have examined the specific neurophysiological mechanisms involved in CLAS, in part because of practical limitations to available tools. To evaluate evidence for possible models of how sound stimulation during brain up-states alters brain activity, we simultaneously recorded electro- and magnetoencephalography in human participants who received auditory stimulation across sleep stages. We conducted a series of analyses that test different models of pathways through which CLAS of slow oscillations may affect widespread neural activity that have been suggested in literature, using spatial information, timing, and phase relationships in the source-localized magnetoencephalography data. The results suggest that auditory information reaches ventral frontal lobe areas via non-lemniscal pathways. From there, a slow oscillation is created and propagated. We demonstrate that while the state of excitability of tissue in auditory cortex and frontal ventral regions shows some synchrony with the EEG-recorded up-states that are commonly used for CLAS, it is the state of ventral frontal regions that is most critical for slow oscillation generation. Our findings advance models of how CLAS leads to enhancement of slow oscillations, sleep spindles, and associated cognitive benefits, and offer insight into how the effectiveness of brain stimulation techniques can be improved.

## Introduction

Closed-loop brain stimulation is a technique in which sensory, magnetic or electric stimulation is timed to modulate endogenous brain activity or states (see Antony et al. (2022) for a discussion of terminology). Its main appeal to neuroscientists is the ability to test relationships between brain activity and function causally, thus adducing strong evidence for the roles of neural events and processes. Closed-loop auditory stimulation (CLAS) of slow oscillations during non-rapid eye movement (NREM) sleep has enjoyed particular attention, perhaps due to the unique (and still somewhat mysterious (Girardeau and Lopes-Dos-Santos, 2021)) physiological processes that take place in sleep.

Sleep is a collection of neurophysiological states with several crucial functions including cleaning waste produced during wake-ful activity (Hauglund et al., 2020), rescaling synapses (Blanco et al., 2015, Timofeev and Chauvette, 2017), and transforming temporarily-stored memories into lasting ones in a process known as memory consolidation (Antony et al., 2019, Diekelmann and Born, 2010, Klinzing et al., 2019, Schabus et al., 2004). These processes seem to rely on sleep-specific neurophysiological events that are most often measured in EEG: slow oscillations, K-complexes (also referred to as ‘N550-P900 complexes’ when evoked by external stimuli), and sleep spindles. Slow oscillations (SOs) refer to low frequency brain activity that is defined variously as *<*1.0, 1.5, 2, or 4 Hz (Carrier et al., 2011, Jaar et al., 2010, Ngo et al., 2013, Vyazovskiy and Harris, 2013). These high-amplitude fluctuations in cortical and subcortical excitability occur during NREM sleep (stages N2 and N3) and appear to be involved in all three processes (Hauglund et al., 2020, Klinzing et al., 2019, Neske, 2016, Timofeev and Chauvette, 2017). N550-P900 complexes are a particular kind of evoked slow oscillation that also occur during stage N2 and N3 sleep (about 550 to 900 ms after sensory input) (Bellesi et al., 2014, Latreille et al., 2020, Riedner et al., 2011). They are highly related to and often considered to be K-complexes, although K-complexes can be generated in the absence of external stimulation. N550-P900 complexes have been linked to sleep continuity and arousal (Halász, 2016). Sleep spindles are transient (*<*2.5 s) 11-16 Hz bursts of activity that are generated through thalamocortical interactions. They frequently co-occur with slow oscillations, may be evoked by sensory stimulation (Sato et al., 2007), and are linked to memory and consolidation processes (see Fernandez and Lüthi (2020) for a comprehensive review).

Closed-loop auditory stimulation of slow oscillations, and subsequent enhancement of memory consolidation of performance on a declarative memory task (e.g. word list learning), was first reported by Ngo et al. (2013). CLAS of SOs has since been repeatedly used in sleep research due to its convenience, ease of use, and temporal precision (see recent reviews Choi et al. (2020), Harrington and Cairney (2021)). As with other forms of brain stimulation, effectiveness can vary considerably across subjects (Nasr et al., 2022). Timing of stimulation seems crucial, and stimulation close to SO peaks (i.e., the ‘up-state’) evokes subsequent SO activity, as well as sleep spindles, and best enhances sleep-related memory consolidation processes (Navarrete et al., 2020). Stimulation during the trough (i.e., the ‘down-state’) can result in a decrease in spindle activity and delta power (Moreira et al., 2021, Ngo et al., 2013). It is thought to be through evoked neural events that memory consolidation is enhanced (Bellesi et al., 2014); however, the neurophysiological mechanisms underlying the effects of CLAS to slow oscillation up-states are poorly understood, and thus it is difficult to optimize and improve stimulation effectiveness. Notably, states are measured most often using a single EEG channel (frontal or central electrode referenced to earlobe or mastoid); this technique captures large-scale, global changes in excitability of cortical tissue. During up-states, neurons are thought to be closer to their firing thresholds, meaning they are more readily activated upon auditory stimulation. However, slow oscillations are not homogeneous phenomena; they originate at different sites and travel along the cortex (Massimini et al., 2004). These dynamics are poorly captured in single-channel EEG. Spatially-resolved techniques such as magnetoencephalography (MEG), which yields information about the relative timing and amplitude of brain responses across brain regions (Baillet, 2017), are needed to explore regional differences in tissue state when stimulation occurs, and to localize its effects.

Evoked brain responses (i.e., event-related potentials in electroencephalograpy; ERPs) are used in sleep research to characterize the evolution of sensory processing according to the depth of sleep (reviewed in Colrain and Campbell (2007)). The amplitude and latency of typical patterns of peaks and troughs, sometimes called ‘components’, are somewhat affected by deepening sleep stage (Niiyama et al., 1994, Ogilvie et al., 1991). The N100 component (i.e., a negative peak occurring about 100 ms after sound onset), is often reported to be attenuated during sleep, whereas P200 (i.e., a positive peak about 200 ms after onset) increases in deeper sleep (Campbell et al., 2010). Some evoked components occurring more than 300 ms after stimulation only appear during sleep (Bastien et al., 2002). Three of these components, labelled N350, N550 and P900 (i.e., negative deflections occurring at about 350 and 550 ms and a positive deflection at about 900 ms post onset) begin to emerge at NREM sleep onset (stage N1) and their amplitude increases with sleep depth (stages N2 and N3). The N550-P900 complex has an asymmetrical shape composed of a trough in which cortical neurons are hyperpolarized, followed by a longer rebound of cortical activity during which neurons are comparatively depolarized and thus more excitable (Latreille et al., 2020). Recent work using electrical source localization models suggests that the spontaneous K-complex, which is similar to the sensory-evoked N550-P900 complex (Halász, 2016), originates in ventral limbic cortex, including medial temporal and caudal orbitofrontal cortex (OFC) (Morgan et al., 2021).

Because the specific auditory pathway involved in closed-loop auditory stimulation is not yet well understood, we must consider the anatomy of the auditory system. There are two main ascending auditory pathways: the lemniscal or primary pathway, and the non-lemniscal or secondary pathway. In the lemniscal pathway, the primary auditory cortex receives projections from the cochlear nucleus via the central nucleus of the inferior colliculus and the ventral division of the medial geniculate body of the thalamus. The non-lemniscal pathway consists of many sets of fibres that send and receive feed-forward and feedback projections widely in midbrain, cortical and limbic areas (Lee, 2015). Of interest here are fibres originating in the brainstem which project to the dorsal division of the medial geniculate body of the thalamus and to secondary auditory cortex, and those which interact with the ascending reticular activating system (ARAS). The ARAS is comprised of many nuclei with connections to the rest of the cortex (Yeo et al., 2013). It also has connections to the locus coeruleus, a group of large nuclei in the brainstem (Bellesi et al., 2014, Poe et al., 2020). The locus coeruleus is the principal source of norepinephrine/noradrenaline in the brain and has widespread (though sparse) projections to the cortex. Both the locus coeruleus and ARAS are involved in transitioning among arousal states, meaning that there is overlap between the auditory non-lemsniscal pathways and the arousal-promoting systems (Bellesi et al., 2014, Wijdicks, 2019).

Evidence suggests that elements of the auditory system, as well as general arousal mechanisms that are independent of the auditory modality (Riedner et al., 2011), are involved in generating the slow oscillations observed in the CLAS effect (Bellesi et al., 2014). However; how, where, and when these systems conspire to produce SOs is unknown, and their contributions to the more commonly-studied cortical EEG evoked responses are not yet understood. Next, we define four possible variations of models of auditory-arousal system interactions, based on the auditory system anatomy described above, ideas proposed in Bellesi et al. (2014), and extent research concerning the origins of evoked and endogenous slow oscillations (e.g., (Halász, 2016, Morgan et al., 2021, Riedner et al., 2011)). Each leads to different predictions in the location, amplitude, and timing of evoked responses to sound in NREM sleep, which can be assessed with temporally- and spatially-resolved neuroimaging methods.

The four models, which vary in the timing and involvement of arousal networks and the location of initial SO generation, are illustrated in Figure 1. In the first model, auditory sensory information reaches the auditory cortex (AC) through lemniscal auditory pathways, where it generates a slow oscillation that propagates locally. In this case, only the auditory cortex should exhibit early auditory evoked components (P100, N100, P200), and in NREM sleep, the evoked slow oscillations should occur earlier and with higher amplitude in auditory cortex as compared with other regions. In the second model, auditory information reaches auditory cortex as above, but then propagates to other cortical regions including frontal areas via structural connections (see Plakke and Romanski (2014)). Upon arrival in the ventral frontal regions, the neural impulse induces a slow oscillation locally, which then propagates across the cortex as previously reported for endogenously-generated SOs (Massimini et al., 2004). In this case, the auditory cortex is not the primary generator of evoked slow oscillations, but the process of SO generation is dependent upon a robust response in the auditory cortex. This model would predict a delayed or absent auditory response in the OFC (orbitofrontal cortex), and that late components would appear first and strongest in ventral frontal regions (like OFC), and then later in other cortical regions. In the third model, as auditory information travels simultaneously through the lemniscal and non-lemniscal pathways, non-lemniscal neurons interact with arousal systems in the brainstem (possibly involving LC) and/or thalamus, which then result in diffuse changes in cortical excitability, leading to the generation of slow oscillations throughout the brain. An early auditory-like response followed by a slower late component should be observed throughout the cortex, although due to the previously-observed tendency for ventral limbic areas (including OFC) to generate endogenous slow oscillations (Massimini et al., 2004, Morgan et al., 2021), we would expect larger-amplitude SOs in the OFC.

**Fig. 1.**
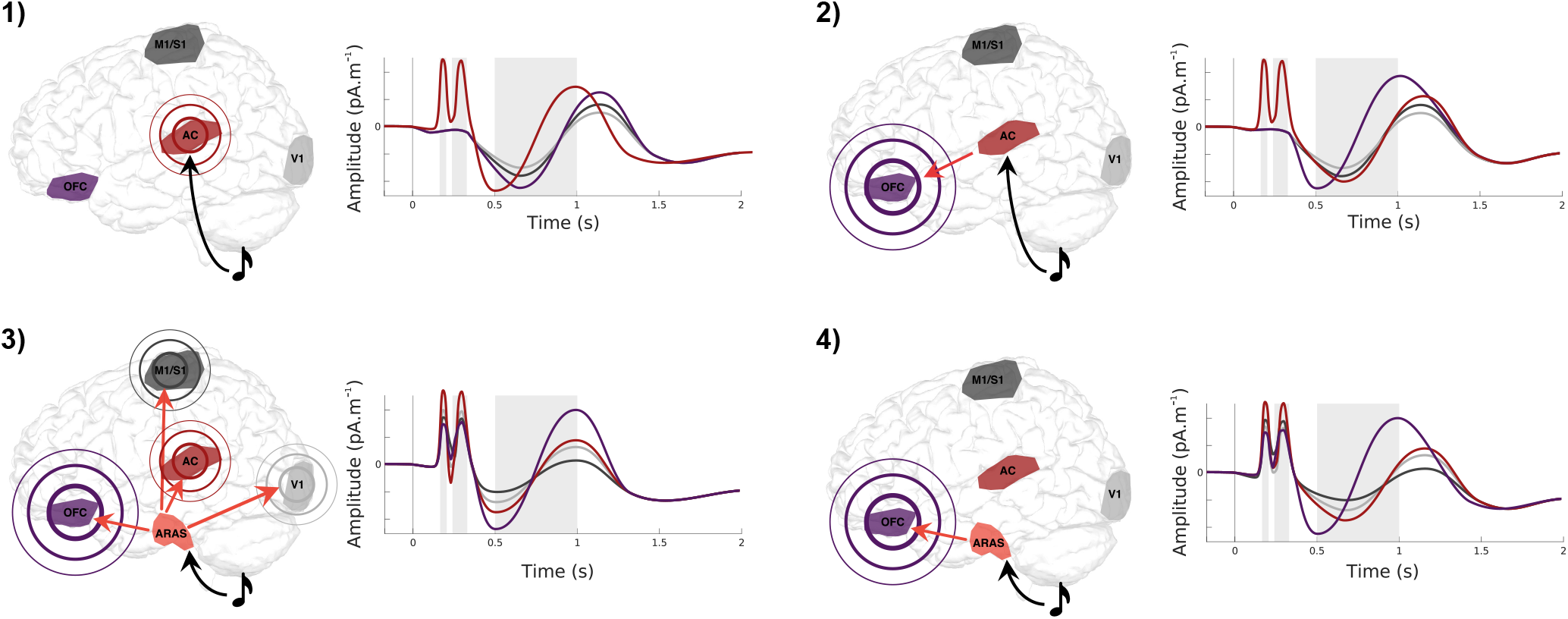
Four models of information flow leading to generation of evoked slow oscillations. 1) SO is evoked by arrival of auditory information in the AC via auditory pathways, from which it propagates. 2) Auditory information arrives in the AC, spreads in cortico-cortical networks to the OFC, which is sensitive to stimulation in sleep and generates SOs, from whence they propagate. 3) The arousal networks in brainstem and thalamus are stimulated, and generate widespread and simultaneous SOs, most strongly in OFC due to its sensitivity. 4) Arousal system is activated as in 3, but SOs are generated first in OFC, from which they propagate to other brain areas. Grey shading indicates the evoked responses components P200, N350 and N550-P900. SO: slow oscillations, AC: auditory cortex, OFC: orbitofrontal cortex. Grey shading indicates time windows of interest that are used in later analyses

In the fourth model, as in the third, auditory information primarily generates the CLAS effect via the non-lemniscal auditory pathway. Contrary to Model 3, in Model 4 the impulse would preferentially stimulate SOs in the OFC, which then propagate to other cortical areas. In Model 3, the SOs would be generated locally (in each region of interest) via common ARAS inputs, whereas in Model 4, the slow OFC activity would induce SOs in other cortical areas. These models can be differentiated using directional functional connectivity metrics in the slow frequency band.

In the present work, we recorded magnetoencephalography (MEG) and EEG data simultaneously in an overnight design, in five healthy young adults, while short, quiet sounds were presented. We first replicate previous results showing that sleep-specific evoked response components emerge in deeper sleep stages. We then take advantage of MEG’s spatial and temporal resolution to explore the distribution of evoked responses over the cortex, their temporal evolution, and their co-occurrence with evoked sleep spindles. We then evaluate evidence in favour of each of these models using MEG. Finally, we explore the relationship of the MEG results to the signal observed in single-channel EEG, as it is most frequently used both to detect SOs and assess the impact of CLAS-SO on evoked oscillations in extent research. Clarifying how sound evokes slow oscillations and sleep spindles has important implications for using CLAS as a causal tool to explore mechanisms of memory, and optimizing its effectiveness in fundamental research and in potential clinical applications.

## Abbreviations

AC: Auditory cortex
ARAS: Ascending reticular activating system
CLAS: Closed-loop auditory stimulation
CN: Cochlear nucleus
EEG: Electroencephalography
ERP: Event-related potentials
FDR: False Discovery Rate
FIR: Finite impulse response
FOOOF: Fitting oscillations & 1/F
IC: Inferior colliculus
LC: Locus cœruleus
LME: Linear mixed effects
MEG: Magnetoencephalography
MGB: Medial geniculate body
NREM: Non-rapid eye movement sleep
OFC: Orbitofrontral cortex
REM: Rapid eye movement sleep
RMS: Root mean square
ROI: Region of interest
SD: Standard deviation
SE: Standard error
SO: Slow oscillation

## Methods

### Participants

Six neurologically healthy young adults without sleep or neurological conditions were recruited from the local community. One dataset was unusable due to a technical problem with the EEG system that precluded scoring. The mean age of the five remaining participants was 21.2 (SD: 1.33; range 19-23), and 3 were female. All subjects reported being in good health with normal hearing, were non-smokers, were not taking medication, had not changed time zones or conducted shift work in the 6 weeks preceding the experiment, and had a normal 6-10 hour sleeping pattern in the three days prior to the experiment (confirmed by sleep log). The night prior to the experiment, participants were asked to stay up one hour later than their habitual bedtime to increase sleep pressure. The day of the experiment they were asked to refrain from consuming caffeine, alcohol, nicotine, and cannabis. Subjects gave written informed consent, and were compensated for their time. The experimental protocol was approved by Concordia University’s Human Research Ethics Committee, and the Research Ethics Board at McGill University. This study was not pre-registered.

### Study design

On the experimental evening, participants first completed a health questionnaire to confirm eligibility; the Munich Chronotype Questionnaire (MCTQ) to document their habitual diurnal rhythms (Roenneberg et al., 2003); and the Pittsburgh Sleep Quality Index (PSQI) (Buysse et al., 1989), which contains 19 self-rated questions which are scored from 0-3 points, where ‘0’ indicates no difficulty, and ‘3’ indicates severe difficulty. The PSQI seven component scores are added to yield a global score between 0 and 21 points. Scores of 5 or more indicates poor sleep quality; the higher the score, the worse the quality. Participants also completed a subjective fatigue scale concerning their current state, which consisted of 7 questions rated on a 5 point Likert-scale. Items on the scale asked about subjects’ levels of tiredness, activity, motivation, interest, concentration, relaxation and general feeling; all ratings were moderate.

Participants were prepared for EEG with electrodes positioned at Cz, C3, and C4; 10-20 system (Homan et al., 1987), noting that the majority of closed-loop studies (whose effects we are trying to better understand) used similar frontal-central electrode positions. To avoid discomfort due to contact with the MEG helmet, occipital electrodes were not used. Participants settled into the MEG scanner in a supine position. A five-minute resting state data acquisition with eyes open was collected, as a contribution to an open-access database (which also houses the current dataset (Niso et al., 2016)). Lights were dimmed to full darkness, and subjects were asked to close their eyes and relax. Sound stimulation was started immediately, with the subject in the awake state, to allow them to get used to the sound and so that the brain response to the wake state could be measured. Standard T1-weighted magnetic resonance anatomical images (1 mm^3^ isotropic voxels) were acquired in a separate session to enable distributed source localization of MEG signals.

### Auditory stimulation

We used a 120 ms synthetic speech syllable (/da/, 10 ms consonant burst, a 30 ms formant transition, and an 80 ms steady-state vowel with a fundamental frequency of 98 Hz), to facilitate comparison of the evoked responses in simultaneously acquired EEG and MEG with previous work (Coffey et al., 2016, 2017, 2021). The stimulus was presented binaurally through Etymotic ER-3A insert earphones with foam tips (Etymotic Research), at 55 dB SPL, which we determined through pilot testing was clearly audible but did not awaken sleeping participants. The stimulus onset interval range was 2800 to 3000 ms (mean *∼*2900 ms), where jitter was selected randomly from a uniform distribution.

One of our main goals is to better understand the neurophysiology behind the CLAS effect, and how it can be optimized. Stimulating only up-states as measured by single-channel EEG would a) limit the total number of stimulations sent, and b) limit our ability to explore the optimal phase of stimulation with respect to excitability state in specific brain locations using MEG, which is not well-captured in the global electrical fields measured using single-channel EEG. We therefore elected to use a continuous sound presentation rather than exclusively stimulating EEG up-states. This design has some potential limitations. The first is that regular sound presentation might limit the depth of sleep achieved. We started presenting sound before sleep, and confirmed that participants were able to fall asleep, and slept adequately, using polysomnography. All participants achieved deep sleep (and even some REM sleep; (see Table 1), suggesting that the auditory stimulation did not impair sleep quality. Second, it is possible that brain responses were reduced due to the frequency and regularity of sound presentation. We included some temporal jitter to reduce predictability, but the inter-stimulus interval is nonetheless kept short in an effort to maximize the number of trials. Bastien et al. found that some components of the evoked responses in sleep are affected by presentation rate (i.e., N350, N550), but not others (P900) (Bastien and Campbell, 1994), suggesting that the slower evoked responses is preserved even with shorter intervals (of around 5 s). We confirmed that we were able to observe these evoked responses across sleep stages using the present inter-stimulus interval (Figure 2). Third, it is possible that brain state at the moment of stimulation is impacted by the previous stimulation, when using short inter-stimulus intervals. In the present study, baseline levels of activity appeared to be restored before 2.5 s following stimulation, implying that processing of new incoming sounds was not strongly influenced by residual activity caused by the preceding sound.

**Table 1.**
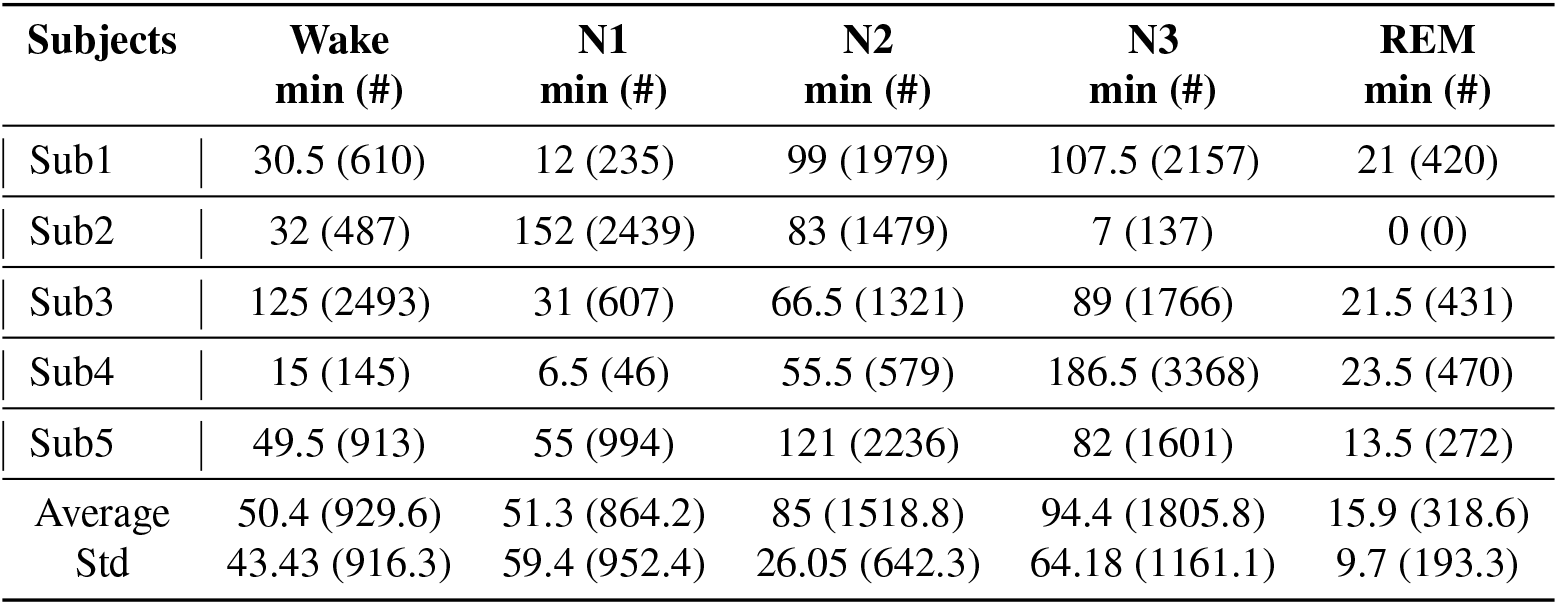
Sleep staging results from usable recordings during which sound stimulation was present, for each subject. Sleep was staged into 30 s epochs, and reported in minutes per stage. The number of auditory stimulations occurring during each stage is reported in parentheses.

| Subjects | Wake<br>min (#) | N1<br>min (#) | N2<br>min (#) | N3<br>min (#) | REM<br>min (#) |
| --- | --- | --- | --- | --- | --- |
| Sub1 | 30.5 (610) | 12 (235) | 99 (1979) | 107.5 (2157) | 21 (420) |
| Sub2 | 32 (487) | 152 (2439) | 83 (1479) | 7 (137) | 0 (0) |
| Sub3 | 125 (2493) | 31 (607) | 66.5 (1321) | 89 (1766) | 21.5 (431) |
| Sub4 | 15 (145) | 6.5 (46) | 55.5 (579) | 186.5 (3368) | 23.5 (470) |
| Sub5 | 49.5 (913) | 55 (994) | 121 (2236) | 82 (1601) | 13.5 (272) |
| Average | 50.4 (929.6) | 51.3 (864.2) | 85 (1518.8) | 94.4 (1805.8) | 15.9 (318.6) |
| Std | 43.43 (916.3) | 59.4 (952.4) | 26.05 (642.3) | 64.18 (1161.1) | 9.7 (193.3) |

**Fig. 2.**
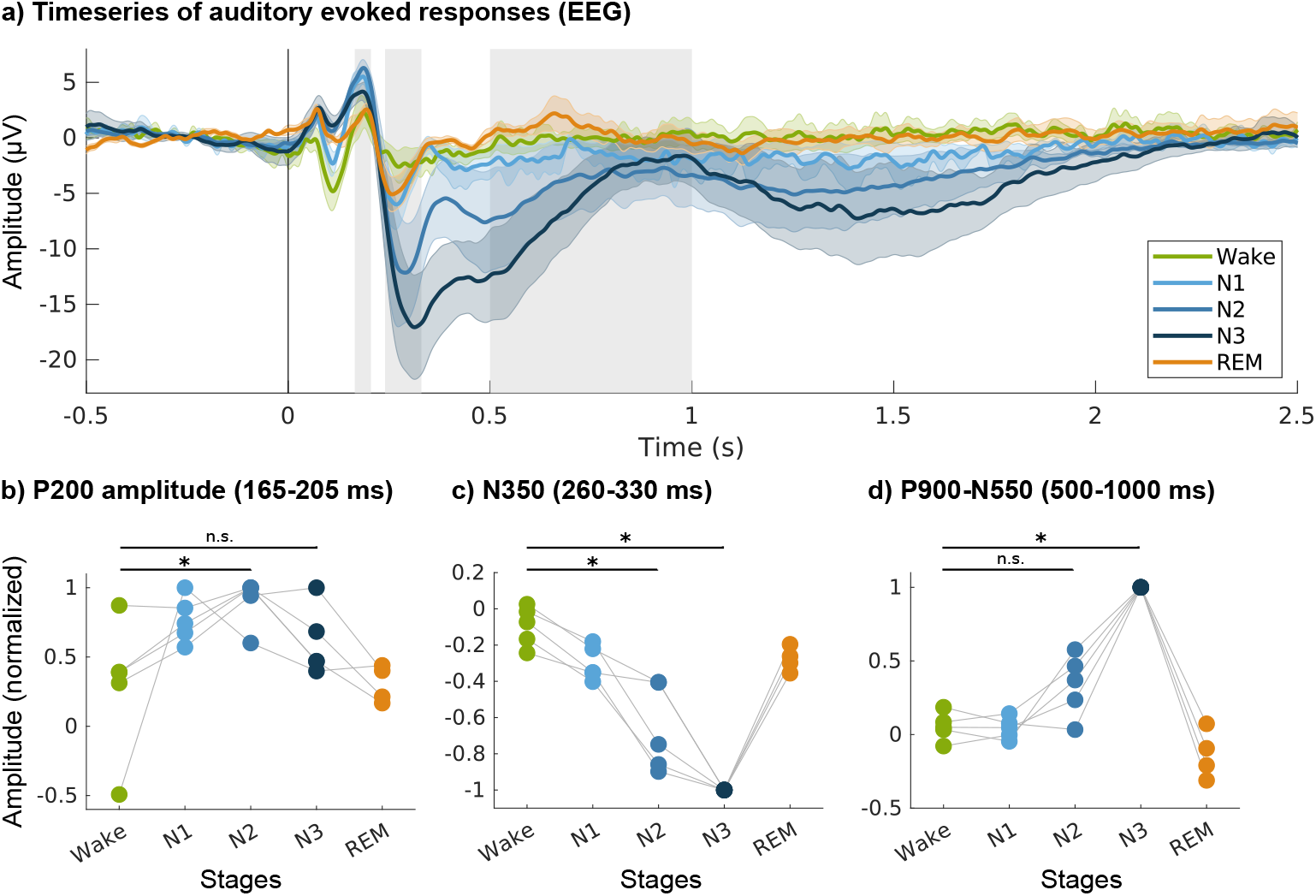
Evoked responses over sleep stages, in EEG (Cz). a) Time series of the auditory evoked responses showing the appearance and evolution of late evoked components (N350, N550, P900) in deeper NREM sleep stages. Coloured lines show means across participants; shaded areas indicate standard error. Grey shading indicates the evoked responses components P200, N350 and N550-P900. b) Amplitudes of the P200 component show little change across sleep stages. Conversely, amplitudes c) at N350 and d) peak-to-trough differences between P900 and N550 are stronger in deeper NREM sleep stages than in Wake. Asterisks denote significant differences between Wake and deeper sleep conditions (Wilcoxon signed-rank test on the means) **\*** p < .05). EEG: electroencephalography; NREM: non-rapid eye movement sleep, REM: rapid eye movement sleep; N1-3: stage 1-3 NREM sleep.

Notably, existing studies that have used closed-loop stimulation and have found memory effects (e.g. Ngo et al. (2013)) often stimulate either pairs or trains of SO peaks, resulting in even shorter inter-stimulus intervals of around *∼*1 s, sometimes with no-stimulation periods of only *∼*2.5 s (Ngo et al., 2013, 2015). For similar reasons, although it is possible that short inter-stimulus interval results in less spindle activity generation due to some stimulations arriving towards the tail end of a spindle refractory period (Antony et al., 2018, Fernandez and Lüthi, 2020), past results with short intervals in a closed-loop setting did successfully generate spindles. We can therefore conclude that the auditory stimulation parameters are compatible with existing CLAS studies and are appropriate for addressing the present questions.

### EEG and MEG data collection

Two hundred and seventy MEG channels (axial gradiometers), five EEG channels (C3, C4, Cz, M1 and M2), two bipolar EMG channels (chin and neck), EOG, ECG, and one audio channel were simultaneously acquired using a CTF MEG System and its in-built EEG system (Omega 275, CTF Systems Inc.). All data were sampled at 2400 Hz, and were recorded in 30-minute ‘runs’. Prior to each run, the head was localized, and head position was stored for use in MEG source localization. Data pre-processing was performed with Brainstorm (Tadel et al., 2011), and using custom scripts (The Mathworks Inc., MA, USA), according to recommended practices (Gross et al., 2013).

### Sleep scoring

Sleep scoring of the data, in which 30 s windows of data are visually inspected and categorized into wake, non-REM sleep stages 1-3, and REM sleep, was accomplished according to AASM practices (Iber, 2007) based on band-pass filtered EEG (0.1-20 Hz, channels ‘C3’, ‘C4’, and ‘Cz’ referenced to left mastoid), EOG (0.1-5 Hz), and EMG (10-58 Hz) channels, which were then down-sampled for ease of handling to 150 Hz. Each epoch was scored by one researcher and confirmed or challenged by a second observer. Discrepancies were resolved via discussion.

### Statistical approach

We elected to study a small number of subjects in an overnight design rather than a larger number of subjects for shorter sleep periods (as is used most nap studies and studies investigating behavioural correlates) so as to be able to capture brain responses across all sleep stages, and to have a very high number of stimulations per subject that hit sleep features of interest (e.g. SO troughs and peaks), or missed them (e.g., stimulations that occurred in the absence of SOs, spindles, and their refractory periods). This design also facilitated an in-depth exploration of phase relationships between EEG and specific brain regions. It is increasingly being recognized that statistical power is dependent upon the amount of data available per subject, particularly in cases in which the within-participant variance is larger than the between-participants variance (Baker et al., 2021). The approach of collecting a large amount of data in a small number of subjects is best suited to research questions that address low-level mechanisms that are likely to be highly similar across participants. It has a long history in non-human primate research, in which research resources and ethical considerations limit sample sizes (please see Baker et al. (2021), Brysbaert and Stevens (2018), Smith and Little (2018) for a discussion and demonstration of the trade-offs between number of subjects and number of observations).

Analysis for basic confirmations of well-established results (e.g., the presence of an evoked N550-P900 complex in NREM sleep stages 2 and 3) were conducted using non-parametric Wilcoxon signed-rank tests on subject means. With our small number of subjects, these results are valid, but are only significant if all five subjects show the same pattern of results. They serve only to confirm that well-replicated phenomena (like the existence of a N550-P900 complex in sleep) are clearly and consistently observed across subjects in this study, as a starting point for addressing the study’s specific objectives. Each subject’s mean values across sleep stages were normalized by dividing by their maximum value, for visualization purposes; this approach was used in Figures 2 and 5.

For the main research questions, we applied linear mixed effects (LME) models (Pinheiro and Bates, 2006), which model the relationship between dependent data and independent data when there is a correlation between observations (e.g., multiple measurements taken from each subject; subject is modelled as a random factor). This class of models takes advantage of the high number of observations per subject (i.e., between 5,000 and 116,000 depending on the research question). LME analyses were conducted in R using the lme4 and emmeans packages (Bates et al., 2015, Lenth, 2022). Alpha values of .05 were used throughout, and FDR corrections for multiple comparisons were applied over the dimensions of interest (e.g., frequency bins) when necessary, using the Benjamini-Hochberg procedure (Thissen et al., 2002). For each linear mixed effects models used in analysis, we visually inspected the histograms and quantile-quantile (Q-Q) plots of the residuals for deviations from normality and homoskedasticity. In models in which clear deviations were present, we excluded outliers by removing values based on thresholds that were defined as 1.5 times the interquartile range (i.e., below Q1 and above Q3; this methods removes data which lie beyond 2.7 *σ* from the mean), and re-ran the models. The use of outlier removal is indicated for each analysis.

### EEG auditory evoked responses across sleep stages

For EEG analysis, we used a ‘Cz’ to left mastoid channel. We first filtered the raw data between 0.1 and 200 Hz (87010-order linear phase FIR (finite impulse response) filter with a Kaiser window and 60 dB stop-band attenuation; the order is estimated using the MATLAB ‘kaiserord’ function, and filter delay is compensated by shifting the sequence to effectively achieve zero phase and zero delay, as per Brainstorm default settings (Tadel et al., 2011)). Data were then down-sampled to 500 Hz, epochs from 1 second pre-stimulus onset to 3 seconds post-stimulus onset were extracted, and the mean during the pre-stimulus baseline period was subtracted from each epoch. Epochs were separated according to the sleep stage during which they occurred, and grouped across an individual’s runs. Subject averages were computed for each sleep stage, a 40 Hz low-pass filter was applied to the average so as to maximize comparability with previous work looking at auditory evoked responses, and mean values from -0.5 s to 0 s were subtracted for better visual alignment across sleep stage time series. Grand average means and standard error were computed for each time series, for visualization purposes.

To quantify the amplitude of the P200 ERP component, we computed average amplitude for each stage between 165 and 205 ms (based on a previously-observed P200 peak in both EEG and MEG to the same stimulus at *∼*185 ms (see Coffey et al. (2017)). The latency of the N350 component appeared to increase in deeper NREM sleep stages (see Figure 2). For Wake, N1, and REM, we used a window of 240 to 280 ms; for N2, the window was defined as 265 to 305 ms, and for N3, 290 to 330 ms after sound presentation. To compute the amplitude of the slowest evoked responses, we first averaged a time window 500 to 600 ms to capture the N550 component, and 800 to 1000 ms for the P900 component. We then subtracted the N550 mean value from the P900 value to get an amplitude measure of the N550-P900 complex. Note that we favour this terminology when referring to activity occurring within this time window to avoid assumptions about whether or not K-complexes are evoked on individual trials. However, ‘P900-N550’ is used in figures to indicate the direction of subtraction in the case the amplitude difference between the two components is measured (i.e., P900 amplitude minus N550 amplitude, which results in a positive peak-to-trough value in case of a successful evoked response).

Later components have broader peaks and are generally less temporally-aligned across subjects, motivating the use of longer windows. We used simple non-parametric Wilcoxon signed-rank tests (one-tailed) to test whether the amplitudes of P200, N550, and N550-P900 complexes differed in N2 and N3 sleep as compared with responses within each time window during wakefulness. We calculated effect sizes for between-condition comparisons used the probability of superiority (PSdep), which is calculated as the number of positive difference scores divided by the total number of paired scores (Grissom and Kim, 2012). Because CLAS is thought to affect memory through the generation of not only slow oscillations, but also sleep spindles nested within them (Ngo et al., 2013), we also investigated changes in spindle band power following sound stimulation, across sleep stages. For each epoch, we computed time-frequency power over a 1-30 Hz frequency band (central frequency: 15 Hz, full-width half-maximum: 0.2 s), using spectral flattening (i.e., power values are multiplied by frequency), in Brainstorm (Tadel et al., 2011). All epochs were averaged by sleep stage within subjects, and a subject grand average difference for N3 minus Wake was computed to define a time and frequency window of interest for further analysis, by visual inspection, which was 0.6-1.2 s after sound onset in a 11-14 Hz frequency band. We then computed the difference between this window and an equivalent window preceding sound offset (-0.6 -0 s after sound onset, 11-14 Hz), for each subject and sleep stage.

It is worth noting that the time window used to measure baseline activity occurs */sim*2s after the onset of the previous sound, which could mean that an incoming sound overlaps with long-range responses to the previous sound. However, as mentioned in the description of auditory stimulation above, baseline levels of EEG activity, including sigma power, seem to be restored */sim*2 s following stimulation making this specific time-window (2.3-2.9 s) adequate for baseline measurement (see (Figure 5b). As above, we used a non-parametric Wilcoxon signed-rank test (one-tailed) to confirm that spindle power amplitude increased across subjects following sound presentation as compared with baseline, specifically in N3 when the effect should be most prominent, and in N2 as a basis for merging of N2 and N3 epochs. These EEG-based analyses serve to select parameters (e.g., time and frequency windows) for further analysis in MEG source space.

In past work, epochs have sometimes been divided into those which produce a strong response (K-complex), and those which do not (e.g., Bastien and Campbell (1992), Dang-Vu et al. (2011)). To explore whether evoked responses are unimodally or bimodally distributed, as would be the case in an all-or-nothing response, we selected stimulations which fell into N2 and N3 sleep stages, yet coincided neither with a slow oscillation nor sleep spindle (as measured in the Cz-to-mastoid EEG channel). We extracted average amplitudes for each epoch during the N550 and P900 windows (as in the analysis above), and computed their difference as a measure of the strength of the N550-P900 complex. We then plotted the distribution of trough-to-peak amplitudes for each subject. Epochs were then sorted into those containing the top 25% and bottom 25% amplitude values. To investigate the relationship between elicited N550-P900 complex and spindle generation, time-frequency plots were computed and averaged for each condition and each subject. Each subject’s mean values across conditions were normalized by dividing their maximum value, for visualization purposes.

### SO detection and phase-binning

We used a Cz-to-mastoid channel to detect slow oscillations using an offline detection algorithm developed by Carrier et al. (2011). In brief, EEG was filtered between 0.16 and 4 Hz and four criteria were evaluated to detect the emergence of a slow oscillation: a peak-to-peak amplitude greater than 75 *µ*V, a negative peak amplitude higher than 40 *µ*V, a duration of the negative peak between 125 and 1500 ms, and a positive peak lasting less than 100 ms (Rosinvil et al., 2021). Detected slow oscillations were then divided into bins representing cortical states.

To explore the impact of up-states and down-states of slow oscillations and question the mechanism of CLAS, 150 ms time windows centred on the detected peak and trough of each oscillation were defined and stimulation occurring approximately simultaneously (defined as occurring within a 150 ms window centred on the detected peak or trough) were sorted accordingly ‘Up-state’ and ‘Down-state’ categories. Stimulation occurring during N2 and N3 NREM sleep stages and outside of a detected slow oscillation (or sleep spindles and their associated refractory periods) was sorted into an additional bin referred to as ‘Clear’. Based on the definition by Antony et al. (2018), the spindle refractory period was defined as a fixed time window lasting 2.5 seconds after the offset of the detected sleep spindle. To maximize the number of stimulations per bin, we combined N2 and N3 NREM sleep stages. On average, the number of stimulations during detected slow oscillations per subject across all phases was 411.8 (SD = 180.7, range = 196-608), and the number of stimulations during the ‘Clear’ condition was 2055.5 (SD = 535.6, range = 1474-2764).

### Spindle detection

Sleep spindles were detected as input to sort stimulations into the Clear condition, such that they coincided neither with a slow oscillation nor with a spindle. Sleep spindles were detected on the Cz (referenced to mastoid) channel, because it is centrally located, near where spindles peak (Cox et al., 2017). Spindles were detected offline using an algorithm which emulates human scoring (Lacourse et al., 2019). In brief, the algorithm filters the EEG signal in the sigma band, 11-16 Hz (as per Lacourse et al. (2019)), and applies four criteria: absolute sigma power (default = 1.25), relative sigma power (default = 1.6), and both correlation (default = 0.6) and covariance (default = 1.3) of the sigma band-passed signal to the original EEG signal. After consultation with the authors, we adjusted the parameters slightly to work better with our EEG data (absolute sigma power = 0.8; relative sigma power = 1.0; covariance = 1.0; correlation = 0.4), and a trained expert confirmed successful detection visually for each subject. As in the analysis of slow oscillations, N2 and N3 sleep stages were merged.

### MEG data processing

Cardiac artifacts were removed from MEG data using Brainstorm’s in-built cardiac detection and source signal projection algorithms (Tesche et al., 1995). Projectors were removed when they captured at least 12% of the signal and the topography of the components matched those of ocular or cardiac origin upon visual inspection. Eye blinks were not detected nor removed, as participants generally had their eyes closed during the recording. Data were filtered between 0.1 and 200 Hz, and down-sampled to 500 Hz. For ERP and ERF analyses, data were further low-pass filtered below 40 Hz, which is common practice to measure the amplitude of evoked auditory response components (e.g., P200).

Auditory event markers were used to define epochs that started 1 s before sound onset and ended 3 s after sound onset as in the EEG analysis. Similar to previous work (Coffey et al., 2021), we used a distributed source-modelling approach. This process estimates activity originating throughout the brain, constrained by spatial priors derived from each subject’s T1-weighted anatomic MRI scan (Baillet et al., 2001, Gross et al., 2013). We imported anatomical data into Brainstorm, and cortical and subcortical (thalamus, hippocampus, and amygdala) structures were combined with the cortex surface (*∼*15,000 vertices). An overlapping-sphere head model was computed for each run; this forward model explains how an electric current flowing in the brain would be recorded at the level of the sensors (Tadel et al., 2011). A noise covariance matrix was computed from 2 min empty-room recordings taken before each session. We computed the minimum-norm estimate (MNE) source distribution model using source orientations constrained to brain structure surfaces, for each run (Hämäläinen, 2009). Source models for each run were averaged within-subject.

### MEG topographies

To explore the topography of the evoked slow oscillations (as indexed by N550-900 complex amplitude), we prepared subject averages in source space for stimulations occurring in Wake, N2, and N3 sleep stages, and extracted time series using the Destrieux brain atlas (Destrieux et al., 2010). As our source models included thalamus, hippocampus, and amygdala in addition to the cortex, we created ROIs for these areas. Because the first two structures are large and aver-aging source-spaced signals over large, elongated or round regions can degrade the signal, we divided them each into three equally-sized ROIs, and visually confirmed that each subject’s divisions were similarly positioned and numbered, thus roughly corresponding across subjects and allowing for meaningful group averages. To maximize the number of trials and thus signal clarity for the sleep analyses, we calculated a weighted average across N2 and N3 sleep stages, proportional to the amount of N2 and N3 sleep collected for each participant (see Table 1). We then created weighted grand averages for each of the Destrieux regions of interest for Wake and combined N2 and N3 sleep (‘N2 & N3’), taking into consideration the proportion of data recorded for individual participants. Each ROI time series was baseline-corrected (-0.5 to 0 s relative to stimulus onset). The amplitude of the N550-P900 complex was computed as described in the EEG analysis (i.e., an average over 500-600 ms was subtracted from an average over 800-1000 ms), and the absolute value was taken. A difference was then computed between the Wake and N2 & N3 conditions. ROIs having outlying values +/- 3SD of the mean across all ROIs was set to +/-3SD, and the values were mapped to colour scales such that red indicates greater activity change relative to pre-stimulation baseline, in sleep). To explore the spatial trajectory of brain activity over time in NREM sleep, we conducted a similar analysis, except that brain activity was computed over six consecutive 500 ms windows starting at -0.5 s relative to stimulation, and the absolute value over each window was displayed on a black-to-red colour scale where red indicates more activity change relative to the pre-stimulus baseline period.

To explore the topography of evoked responses in the spindle range, we computed spindle band power as in the EEG window in Wake and NREM (N2&N3) for each region of interest, and mapped the power difference to a blue-red scale where red indicates more spindle activity in sleep. These average activity topographies were used to present spatial information concerning evoked responses qualitatively, and as input to confirm and select regions of interest for further investigation of evoked responses in source space.

For a visual comparison of the topography of the evoked slow responses and spindle band activity, we used the FOOOF al-gorithm (https://fooof-tools.github.io/fooof/) as implemented in Brainstorm (frequency range of analysis: 1-40 Hz; otherwise using default parameters: peak width limits: 0.5-12 Hz, maximum number of peaks: 3, min peak height = 3 dB, proximity threshold: 2 standard deviations) to separate oscillatory from 1/f (frequency) brain activity (Donoghue et al., 2020), and averaged peak values for slow oscillations (0.5-1.5 Hz) and spindles (11-17 Hz) separately. We used the wider spindle frequency range here for best comparison with prior literature, although it is slightly different than the 11-14 Hz range selected based on the evoked spindle power in our own dataset. Note that due to the study design, brain activity captured in this analysis includes a combination of evoked and endogenous activity, meaning that we limit our analysis to a qualitative observation of similarities; a specialized study design that clearly separates evoked from endogenous oscillations would be needed to quantitatively investigated questions of similarity more thoroughly. Note that endogenous slow oscillations and stimulus-evoked K-complexes share many similarities and there is currently no normative data for discriminating evoked responses from other slow waves (Halász, 2016).

### MEG time series analysis

Based on the models that we propose to evaluate (Figure 1) as well as the topographical analysis showed in Figure 4, five bilateral cortical regions of interest (ROIs) based on the Destrieux atlas (Destrieux et al., 2010) were defined in subject space for each participant. The selected ROIs are based on: ‘pre-central gyrus’, ‘post central gyrus’, ‘occipital lobe’, and ‘sub-callosal gyrus’, and the ‘transverse temporal’ and ‘planum temporale’ gyri, which were merged to capture the auditory cortex. Each ROI was defined in each hemisphere. The size of each ROI was manipulated by progressively increasing or decreasing ROI area, so as to be similar across ROIs and hemispheres (across all ROIs, all subjects, mean area: 10.3 *cm*^2^, SD: 0.2 *cm*^2^, range: 9.99 to 10.88 *cm*^2^). We preferred to use ROIs of similar size, as we extract the mean activity from each ROI. In general, we kept left and right hemisphere homologue ROIs separate during the analyses so as to explore the possibility of hemispheric differences, for which there is some limited support. For example, Achermann et al. (2001), Morgan et al. (2021) found that slow oscillatory activity was stronger in left than right ventral limbic areas, using EEG.

For MEG analysis investigating the contribution of specific ROIs, we sorted epochs (i.e., -1 to +3 s relative to sound onset) by sleep stage, and produced evoked response plots, as in the EEG analysis (see Supplementary Figure 1). We have not included SNR measurements, because they are confounded by the nature of the evoked design. That is, if we measure an evoked response we can be confident that we have adequate SNR to measure it, but if we do not, we cannot clearly distinguish between a region not producing a response and our inability to pick it up due to insensitivity (note that we compare results within-region whenever possible). To evaluate the presence or absence of specific peaks in the evoked response, we then measured the mean amplitude over a window centered on P200 in EEG (165 and 205 ms post stimulation) as well as the difference in amplitude within a time window capturing the N550-P900 slope as observed in EEG ERPs (i.e., the average amplitude over 500ms to 600 ms subtracted from the average amplitude from 800ms to 1000ms) for each run and for each ROI. Extracted values for each epoch were then used in LME statistical models to assess the impact of sleep stage on responses elicited by sound across sleep stages.

### Directed phase-transfer entropy

Functional connectivity between the AC and OFC was estimated using directed phase-transfer entropy (Lobier et al., 2014), in N2 and N3 sleep only, as these contain the evoked responses of interest (i.e., SOs and sleep spindles). In brief, the time series is described by its instantaneous phase, in a similar fashion to Wiener-Granger causality (Hillebrand et al., 2016).

We used common frequency band divisions: ‘delta’, ‘theta’, ‘alpha’, ‘beta’ and ‘low gamma’, with the additional separation of 0.5*−*1.5 Hz and 11*−*17 Hz as ‘slow oscillation’ and spindle or ‘sigma’ bands, due to their relevance to the research questions, and because there is some evidence that very slow oscillations (*<*1.5) are somewhat distinct in characteristics and function from 2–4 Hz oscillations (Brodt et al., 2023, Kim et al., 2019, Steriade et al., 1993). Note that while slow oscillations within the 0.1 to 0.5 Hz range have been included in our slow oscillation analyses elsewhere, for example to detect slow oscillations or visualize time-locked evoked responses, the 4 s epochs used for the connectivity analysis only allow us to characterize frequencies above about 0.25 Hz (i.e., 1*/*4 s); we therefore raised the filter to 0.5 Hz. In sum, signals were band-pass filtered in the following frequency bands: 0.5*−*1.5, 2*−*4, 5*−*7, 8*−*10, 11*−*17, 18*−*29, and 30*−*58 Hz.

Directed phase-transfer entropy was computed for each pair of ROIs within each frequency band, and normalized between -0.5 and 0.5. The sign of the result indicates the dominant direction of functional connectivity. Directed phase-transfer entropy results for each trial were then entered in LME statistical models.

### Up-state and down-state detection in EEG and MEG

We explored the role of tissue state in specific regions and its relationship to EEG by detecting up and down-states in extracted time series from each ROI over each *∼*30 min recording. We used the same detection algorithm for the MEG data (extracted from regions of interest in source space) as had been used for EEG (Cz) signals, to maximize comparability. We first isolated periods of sleep that had been manually scored as N2 or N3, and filtered the data in the slow wave frequency band (0.1 to 2 Hz). Troughs and peaks were automatically detected (using the ‘findpeaks’ function implemented in MATLAB), and the amplitude difference between peak and trough amplitude was computed for each detected peak. Trials were sorted by amplitude differences, and epochs containing the top 25% of peak-trough amplitude for each subject were selected as our Up-state condition for further analyses. Similar processing pipeline was used to select troughs (Down-states) by detecting peaks after inverting the signal (i.e., multiplication by -1).

Epochs were defined around the detected peak (Up-state) in the EEG signal (-2 to +2 s) to examine the correlation between the EEG signal and MEG signal extracted from ROIs. Correlation of the signal from the EEG sensor Cz and each extracted signal was then computed and the correlational relationship between signals was investigated using LMEs. The instantaneous phase of the signal in EEG as well as in each ROI was extracted using the Hilbert transform of the signal.

Finally, to investigate the role of cortical tissue excitation state on EEG brain evoked responses, incoming sounds were sorted according to whether they coincided with a local up-state or down-state in each ROI (as defined by a 150 ms window centred on the detected peak or trough), and as before, 4 s epochs (-1 s pre-stimulation onset to 3 s post-stimulation onset) were defined. The elicited N550-P900 complex amplitude was measured as previously described and the sigma power post-stimulation was calculated using the root mean square (RMS) value of the signal filtered between 11 and 14 Hz, 0.6 to 1.2 s after stimulation from which a pre-stimulus value (-0.6 to 0 s) was subtracted to emphasize sound-evoked spindle band power, as in previous analyses.

## Results

### Sleep

Although sleep quality was not optimal, all subjects were able to sleep sufficiently in the physically-restricted and unfamiliar scanner environment so as to have a large number of trials (see Table 1). Sound stimulation was started before sleep onset and all subjects were able to fall asleep, but as sleep pressure wore off, participants sometimes reported that when they awakened, the stimulation distracted them from falling asleep again. Participants could ask for the stimulation to be turned off until they were once again asleep. Two of the five participants availed themselves of this option. Once they were sleeping, turning on the sounds did not re-awaken them. Unstimulated periods were not included in the analyses and 30 s epochs containing movement were discarded. On average, participants had 246.6 mins of movement-free, stimulated recordings in sleep (including NREM and REM sleep, but not Wake; SD = 26.6 mins). Every subject, except one (‘Sub2’), had *>*10 mins of N3 and REM sleep. See Table 1 for details. In analyses which combined N2 and N3, weighted averages within-subject were used to account for the differences in sleep stage duration. As the study concerns physiological rather than behavioural research questions, these results are sufficient to investigate auditory brain responses during sleep.

### Effect of sleep stage on brain responses evoked by auditory stimulation

#### Auditory evoked responses in EEG

To confirm that the study design and amount of sleep stimulations acquired across sleep stage were adequate to observe previously reported state-dependent differences in evoked responses, we analyzed evoked responses across brain states in EEG. The evoked responses for each observed sleep stage in EEG, averaged across subjects, are presented in Figure 2a. Qualitatively, the early evoked responses that are associated with auditory processing in wake states (i.e., P100, N100, P200) do not appear to be strongly influenced by sleep stage, whereas the later components (i.e, N350, N550, P900) that appear only in sleep are stronger in deeper NREM stages.

We used simple non-parametric Wilcoxon signed-rank tests (one-tailed) to evaluate whether evoked response amplitudes differed between wakefulness and deeper NREM sleep stages (N2 & N3), which are the brain states in which CLAS of SOs is normally used. P200, as a representative measure of the classical auditory evoked response, was significantly greater in N2 sleep as compared to wake (Z = 15, p = .03, PSdep = 1.0), but the Wake vs. N3 difference was not significant (Z = 3, p = 0.31, PSdep = 0.8). The N350 component, instead, was significantly larger in NREM sleep stages 2 and 3 (Z = 15, p = .03, PSdep = 1.0 for both). The N550-P900 complex (i.e., P900 - N550 amplitude) was significantly greater in N3 sleep than Wake (Z = 15, p = .03, PSdep = 1.0), but did not reach significance in N2 sleep (Z = 14, p = .06, PSdep = 0.8). These results are used to confirm that the slow evoked responses that emerge in deeper sleep (N2 and N3) and have been previously reported (see Bastien et al. (2002), Colrain and Campbell (2007)) are present in our sample, as a starting point for spatially-resolved MEG analyses. To increase the number of trials for subsequent analyses, we combine N2 and N3 sleep stages, as N350 and N550-P900 components are observed in both stages (Colrain and Campbell, 2007) (although noting that the N2 effects are smaller than in deeper, N3 sleep). Importantly for the purposes of using LME models, the number of observations included in the statistical analyses for the main questions concerning the effects of sound stimulation in N2 and N3 sleep are over 16,000 per region of interest (see Table 1).

#### Slow oscillations evoked by sound in sleep in MEG source space

To confirm that the orbitofrontal regions of interest is involved in the evoked response to sound, we computed the topography of evoked activity differences between Wake and N2 & N3, displayed in Figure 3a. Widespread activity differences are observed in ventral and orbital frontal regions (blue arrows), and inferior frontal gyrus (e.g., Destrieux regions ’G_subcallosal L’, ’Lat_Fis-ant-Vertical L’, ’G_cingul-Post-ventral L’, ’G_front_inf-Orbital R’, ’S_circular_insula_ant L’). On the basis of these observations, we selected Destrieux atlas regions (‘G_subcallosal’ left and right) as representative of ventral regions exemplifying the SO evoked response (see Methods for details, and Figure 6b for a visualization of the resulting ROI). For visual comparison only, we plotted topographies of slow frequency oscillatory activity during N2 & N3 stages (using the FOOOF algorithm) (Figure 3b). In general, we observed some overlap between the evoked N550-P900 complex topography (Figure 3a) and the slow oscillation activity (Figure 3b). There was more SO activity in dorsolateral frontal regions in b; however, as noted in methods, the specific study design does not allow us to clearly separate (and therefore quantitatively compare) evoked vs. endogenous SO topographies. We also take advantage of MEG’s temporal resolution to visualize the time course of the evoked responses, see Figure 4. In the earlier time window (0 to 0.5 s), the auditory cortex is bilaterally active (blue arrow). In later periods (e.g., 1.5 to 2.0 s), the orbitofrontal regions show greater change with respect to pre-stimulus baseline. The auditory regions also show activity changes in later periods. Note that the P900 component is not prominent in this representation, which displays difference from baseline, because as shown in the time series average (Figure 4 (bottom)), amplitude peaks close to the pre-stimulus level (at 0.9 s) before once again decreasing (at 1.5 s). We nonetheless see activity in the OFC related to the late component of the EEG response around 1.5-2 s (blue arrow). The evoked response returns to baseline *∼*2-2.5 s post stimulation, which is prior to the next stimulation (*∼*2.9 s).

**Fig. 3.**
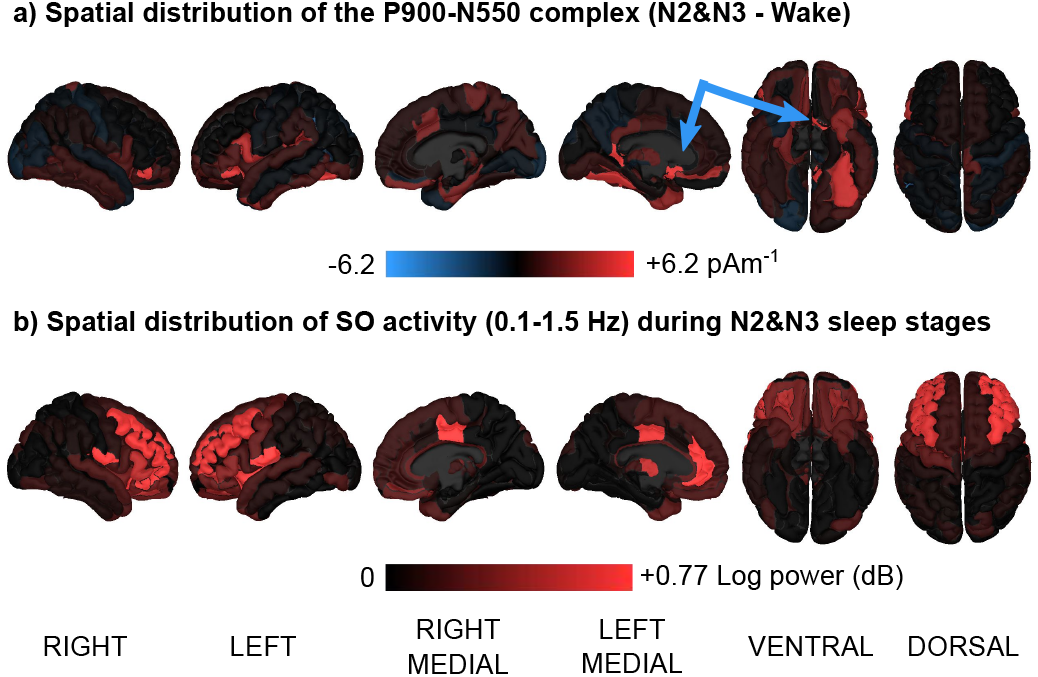
Topographies of a) evoked activity differences between wake and NREM sleep (stages N2 and N3 combined). Red indicates more activity change relative to pre-stimulus baseline during the auditory-evoked N550-P900 complex in sleep as compared to wake. The evoked N550-P900 complex is specific to sleep and shows widespread activity change in ventral and orbital frontal areas (see also Figure 2b,d for topographies of evoked activity in NREM sleep over time). b) Topography of slow frequency oscillatory activity during N2 & N3 sleep stages (as indexed using FOOOF), for reference; note that due to the study design, both spontaneous and evoked slow oscillations are represented. Blue arrows indicate activity in the orbitofrontal cortex. FOOOF: fitting oscillations & 1/F; SO: slow oscillation; N2-3: stage 2-3 NREM sleep.

**Fig. 4.**
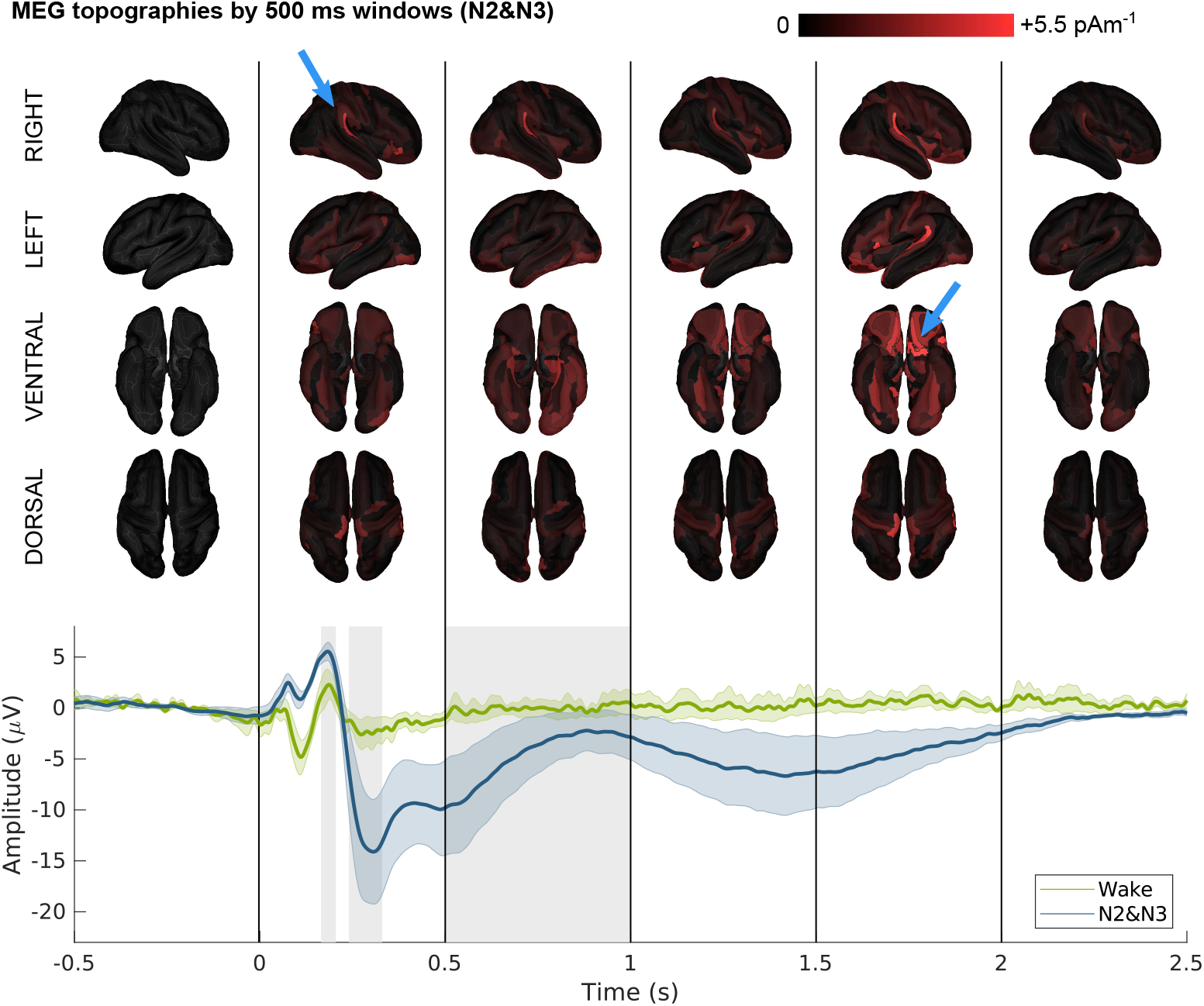
Topographies of evoked activity in NREM sleep over time (stages N2 & N3 are combined). Brighter red indicates relatively more activity change relative to pre-stimulus baseline. The EEG time series of evoked responses for Wake and N2&N3 is shown (bottom) for reference. Coloured lines show means across participants; shaded areas ndicate standard error. Grey shading indicates the evoked responses components P200, N350 and N550-P900. Blue arrows note where the activation occurs. Grey shading ndicates the evoked responses components P200, N350 and N550-P900. MEG : magnetoencephalography;EEG: electroencephalography; NREM: non-rapid eye movement sleep; N2-3: stage 2-3 NREM sleep.

**Fig. 5.**
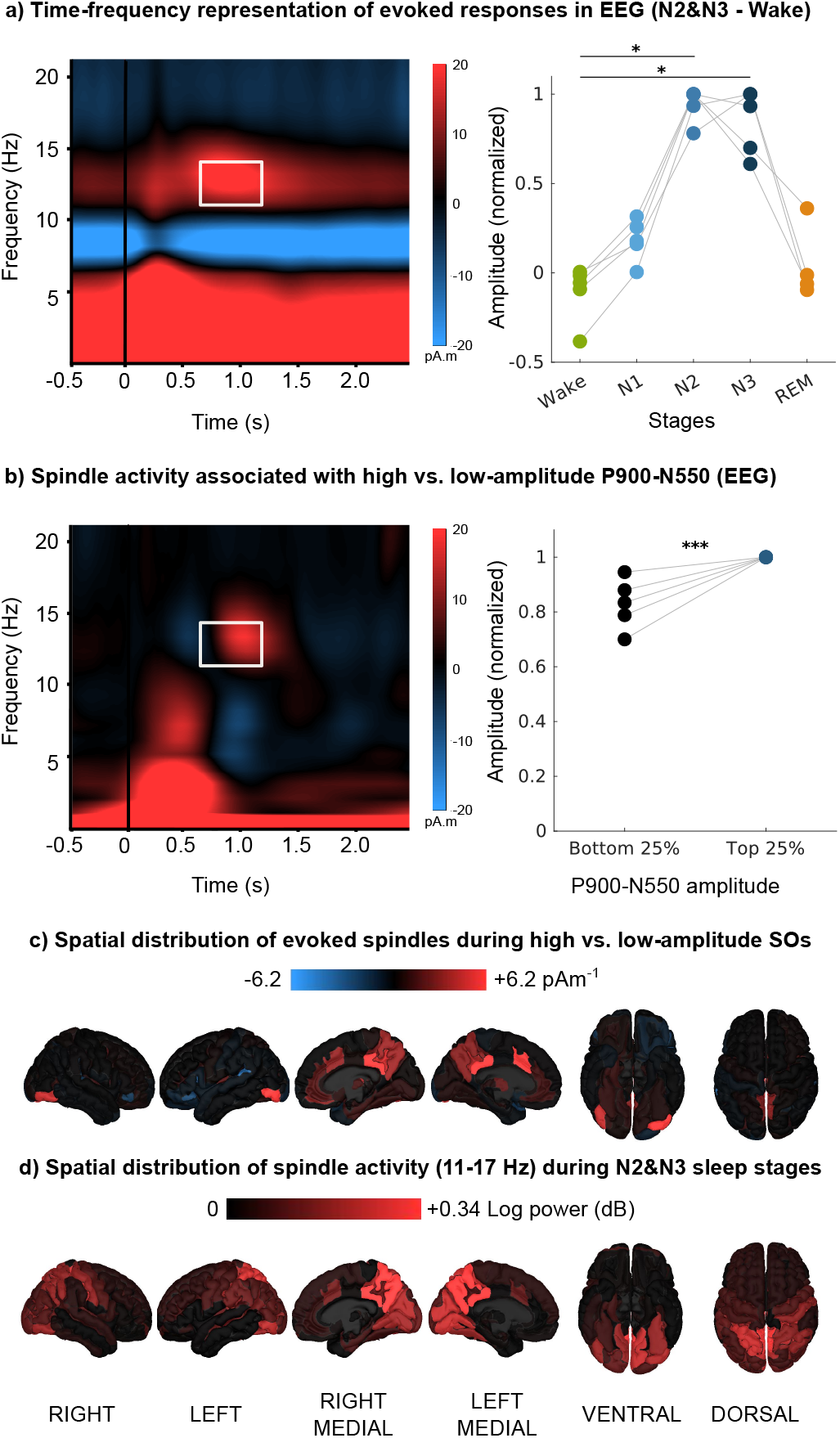
Sound-evoked sleep spindles in EEG (Cz). a) The time-frequency plot averaged across all subjects (left) shows increased spindle power in N2 & N3 sleep 0.6 to 1.2 s after sound onset in a frequency band of 11 to 14 Hz (white rectangle). Spindle power in that frequency range as compared with spindle power in the pre-stimulus baseline (-0.6 to 0 ms) is shown at right. Each subject is represented by its mean. Asterisks denote significant differences between Wake and N2 & N3 (**\*** p < .05). b) The difference time-frequency plot averaged across all subjects (left) comparing spindle activity when epochs are divided according to the strength of the N550-P900 complexes that they produce (i.e., Top minus Bottom 25%). Stronger spindle power is observed to be coincident with stronger N550-P900 complexes in the EEG time-frequency plot, suggesting that spindle activity is positively related to evoked N550-P900 complex amplitude. Extracted values (white rectangle shown on left) of spindle power in the Top 25% are higher across subjects (right).Each subject is represented by its mean. Asterisks indicate a significant difference between the Top vs. Bottom 25% (**\*\*\*** p < .001). c) The topography of spindle power differences (in MEG) according to evoked N550-P900 complex strength (Top vs. Bottom 25% of N550-P900 complexes averaged across subjects) shows ncreased activity in thalamus and frontal midline structures coincident with stronger N550-P900 complexes. d) The topography of spindle power in the 11-17 Hz range across all N2 & N3 sleep epochs, for comparison; note that due to the study design, both spontaneous and evoked sleep spindles are represented. EEG: electroencephalography; N2-3: stage 2-3 non-rapid eye movement sleep 2 and 3, SO: slow oscillations.

To better characterize the sound-evoked N550-P900 complex, we investigated whether it represents an ‘all-or-none’ response, as suggested by some previous work in which epochs have been separated according to the presence or absence of an elicited N550-P900 complex (e.g., Bastien and Campbell (1992), Dang-Vu et al. (2011), Laurino et al. (2014)). In closed-loop auditory stimulation applications, up-states are preferentially stimulated as they produce the greatest enhancement in slow oscillations and evoked spindle power (Ngo et al., 2013). In the current analysis, however, we focused on periods of time in which sound presentation did *not* coincide with slow oscillations or sleep spindles (in EEG), to observe the evolution of the evoked response in the absence of endogenous sleep-related events (later sections explore the effect of cortical tissue excitability on the EEG evoked response).

We first identified trials that occurred within spindle and SO-free stage N2 & N3 sleep (as identified by automatic detection algorithms). The mean number of trials included per participant was 2284.2 (SD = 650; individual totals: Sub1 = 2762, Sub2 = 1325, Sub3 = 1910, Sub4 = 2850, Sub5 = 2574). In general, each subject’s N550-P900 complex amplitude distribution was centred around zero (means (SD) by subject: Sub1: 4.76 *µ*V (43.2), Sub2 = -.04 *µ*V (22.1), Sub3 = 4.54 *µ*V (42.5), Sub4 = 14.89 *µ*V (54.3), Sub5 = 4.26 *µ*V (37.7)). Interestingly, we do not observe a bimodal distribution that might suggest that sound produces a clear evoked slow oscillation in a binary fashion (see Supplementary Figure 2). This analysis does not bear directly on our main research questions concerning the information flow leading to generation of evoked slow oscillations, but rather contributes to an ongoing discussion of the nature of ‘successful’ sound stimulation, and the optimization of the CLAS technique. We further investigate the relationship between evoked N550-P900 amplitude and spindle-band activity in the next section, using the present analysis to separate trials into those producing relatively higher amplitude N550-P900 complexes (i.e., top 25%) and those which had lower values during the same time window (i.e., bottom 25%).

#### Sleep spindles evoked by sound in sleep

To evaluate differences in how sound evokes sleep spindles across stages of consciousness, we calculated mean time-frequency plots for Wake and N2 & N3 across all subjects and epochs, and computed their difference, to identify a time and frequency window for further analysis (see Figure 5a). Evoked responses were maximally different between Wake and N2 & N3 starting about 0.6 s and ending about 1.2 s after sound onset, and within a 11-14 Hz frequency band, by visual inspection. All subjects showed greater spindle power in N2 and N3 than Wake using simple non-parametric Wilcoxon signed-rank tests (one-tailed) across subject means (Z = 15, p = .031, PSdep = 1.0 in both cases; see Figure 5a (right)).

To more robustly evaluate differences in evoked spindle power as a function of the N550-P900 complex’s amplitude, we conducted LME analyses at the single trial level, using subjects as a random effect. We removed outliers from each Condition (Top 25%, Bottom 25%) by excluding values based on thresholds defined as 1.5 times the interquartile range (i.e., below Q1 and above Q3). The mean percentage of retained epochs across subjects was 95.8% (SD = 4.0). We compared a model with N550-P900 complex’s amplitude Condition (either Top 25% or Bottom 25%) as a fixed effect to a null intercept-only model and found that the addition of Condition significantly increased model fit (*χ*^2^(1) = 67.04, p < .0001). The intercept estimates the mean spindle power increase post-stimulation generating low amplitude N550-P900 complexes (Bottom 25% condition). There was a significant main effect of Condition, indicating statistically stronger spindle power in the Top 25% of epochs than in the Bottom 25% of epochs (*β* = 3.83e-01, SE = 4.66e-02, t(5454) = 8.21, p *<* .0001) (see Figure 5b).

The topography of evoked spindle activity differences between Top vs. Bottom 25% of evoked N550-P900 complexes is displayed in Figure 5c. Increased activity was observed mainly in medial regions (e.g., Destrieux regions ‘G_cingul-Post-dorsal’, ‘G_and_S_cingul-Mid-Ant’, ‘G_cingul-Post-dorsal, ‘S_subparietal’), and also in lateral occipital regions (e.g., ‘G_and_S_occipital_inf’). An increase in thalamic activity is also observed. For visual comparison only, we plotted topographies of spindle-band (11-17 Hz) oscillatory activity during N2 & N3 stages (using FOOOF). In general, we observed considerable overlap between the evoked spindle topography as described above (Figure 5c) and activity measured in the spindle band across N2 and N3 sleep stages using the FOOOF algorithm (Figure 5d), with the strongest activity observed in medial regions. As noted in Methods, the study design does not allow us to cleanly separate and therefore quantitatively compare evoked vs. endogenous spindle topographies. These analyses take advantage of our MEG approach to inform us about the spatial distribution of the co-occurrence of the strongest evoked SOs and coupled spindles.

#### Temporal evolution of auditory evoked responses in MEG source space

To address the main research questions evaluating evidence for slow oscillations generation models, we first explore the time course of neural events in specific brain regions to compare activity within cortical regions between sleep and wake states. We extracted time series from cortical regions thought to be involved in the generation of cortical auditory evoked responses (AC and OFC) as well as other sensory regions used as controls (S1/M1 and V1). We averaged the extracted signals across trials and hemispheres according to sleep stage and measured the amplitude of activity during specific time windows (P200 and the N550-P900 complex), for each subject (see Figure 6). We observed peaks of activity in the AC and OFC at about 100 and 200 ms after sound onset (P100, P200), with OFC peaking slightly later than AC, and troughs in OFC and AC occurring around 350-550 ms post-onset (N350, N550). There is some evidence of brain activity in V1, M1 and S1 within 300 ms of sound onset, but peaks appeared delayed or indistinct with respect to AC and the EEG timeseries (in Figure 2).

**Fig. 6.**
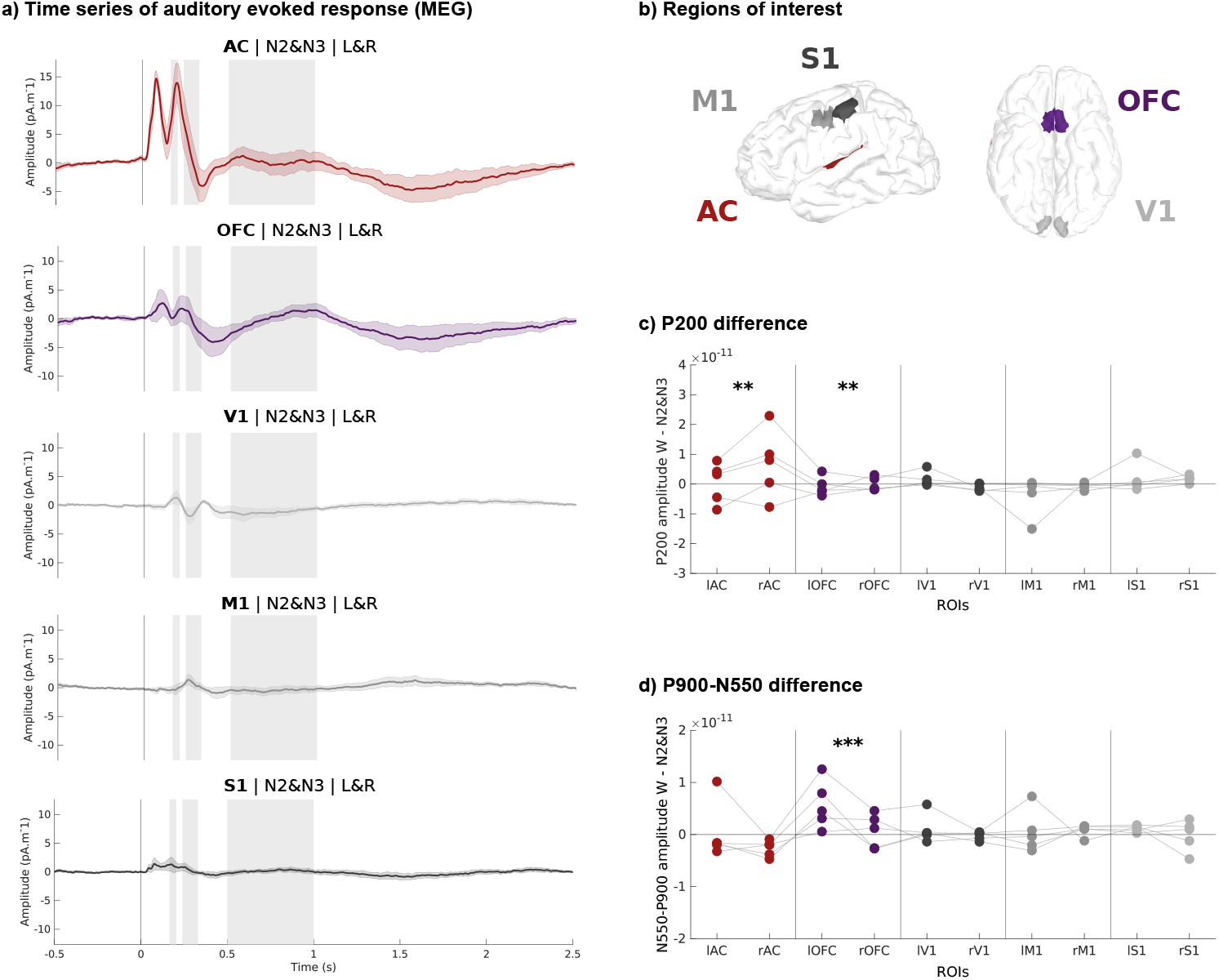
Evoked responses in N2 & N3. a) Time series, averaged across subjects and hemispheres, for each region of interest. Coloured lines show means across participants; shaded areas indicate standard error. Grey shading indicates the evoked responses components P200, N350 and N550-P900. b) Schematic view of regions of interest. P200 amplitude differences between Wake and N2 & N3 for each region of interest and each hemisphere. Each subject is represented by its mean. Asterisks denote significant differences between Wake and N2 & N3 (**\*\*** p < .01) d) N550-P900 complex amplitude differences between Wake and N2 & N3 for each region of interest and each hemisphere. Each subject is represented by its mean. Asterisks denote significant differences between Wake and N2 & N3 (**\*\*\*** p < .001). MEG: magnetoencephalography; N2-N3: non-rapid eye movement sleep stages 2 and 3; L&R: Left and Right brain hemisphere homologues

We then conducted LME analyses on P200 amplitude, using subjects as a random effect to compare amplitude between Wake and N2&N3. We removed outliers from each ROI by excluding values based on thresholds defined as 1.5 times the interquartile range (i.e., below Q1 and above Q3). The mean percentage of retained epochs across subjects was 90.4% (SD = 5.90). Using a null-intercept linear mixed effect model we found that only AC (*β* = 10.14, SE = 0.48, z(5.63) = 21.14, p < .0001) and the OFC (*β* = 2.81, SE = 0.48, z(5.6707) = 5.84, p = .001) across Condition (Sleep and Wake merged) are statistically greater than zero, as suggested by Figure 6. This analysis shows that auditory information is present at P200 across sleep and wake states only in the OFC and AC regions of interest, and motivates focusing on them to assess the impact of sleep state (Condition).

Using a new linear mixed effects model with ROI as fixed effect and Subject as random effect (Amplitude *∼* 1+ ROI +(1|Subject)), we found a main effect of ROI, meaning that both AC and OFC are significantly different across the sleep and wake Conditions (F(1,76138) = 832.77, p < .0001). Adding the interaction between Condition and ROI improved model fit (Amplitude *∼* 1+ ROI + Condition + ROI * Condition + (1|Subject)) as compared with fixed effects of ROI and Condition only (*χ*^2^(1) = 29.87, p < .0001). We do not find a main effect of Condition (Wake, Sleep on P200 amplitude across ROIs (F(1, 72083) = 0.18, p = 0.67) but there is a main effect of ROI (F(1, 76138) = 832.77, p < .0001) as in the ROI-only model, with AC estimated mean amplitude (M = 10.17, SE = 1.08) being higher than OFC estimated mean amplitude (M = 2.78, SE = 1.08). There is a main effect of interaction between ROI and Condition (F(1, 76134) = 29.46, p < .0001), meaning that the ROIs do not behave similarly across wake and sleep states. Post-hoc analysis using estimated means showed that AC amplitude in Sleep is lower than in Wake (Sleep-Wake difference, M = -1.80, SE = 0.44, p = .0003) and that OFC amplitude is higher in Sleep than Wake (Sleep-Wake difference, M = 1.52, SE = 0.46, p = .0052).

This pattern of results shows that auditory information is present in the AC and OFC in both wake and sleep states. However, these regions are differently impacted by sleep state, with AC showing smaller P200 responses and OFC showing larger P200 responses in NREM sleep as compared with wake. This result is most consistent with Models 3 and 4, in which auditory information coming from the non-lemniscal pathway is leading to changes in excitability of tissues within the OFC. At this stage we cannot rule out Models 1 and 2, because feed-forward information via the lemniscal pathway could subsequently reach OFC from AC, albeit with a neural conduction delay. However, diminished P200 amplitude in the AC during sleep suggests a lower likelihood of the lemniscal pathway through AC playing the key role in SO generation in OFC in sleep.

In an exploratory basis, we also tested Hemisphere (Left and Right) as a fixed effect, as compared with the null model (*χ*^2^(1) = 71.49, p < .0001). The results suggest that on average across Condition and ROI, P200 amplitude is greater in the left hemisphere than in the right hemisphere (Left-Right difference, M = 2.17, SE = 0.26, p < .0001).

Next, we conducted LME analyses to investigate the elicited N550-P900 complex amplitude, testing our hypothesis that change in amplitude would be greater in Sleep than in Wake (as seen in the time series average, see Supplementary Figure 1). We removed outliers from each ROI using the same process as defined in the P200 analysis. The mean percentage of retained epochs across subjects was 89.4% (SD = 6.18). First, using a null-intercept linear mixed effect model we found that only the OFC across Condition (Sleep and Wake merged) is statistically greater than 0 (*β* = 1.15, SE = 0.26, z(8.1545) = 4.5, p = .002).

Focusing on the OFC only (Amplitude *∼* 1 + Condition +(1|Subject)), we found a main effect of Condition (F(1, 3946.3) = 20.7, p < .0001). Post-hoc analysis using estimated means showed that OFC N550-P900 complex amplitude in Sleep is higher than in Wake (Sleep-Wake difference, M = 2.17, SE = 0.48, p < .0001).

The absence of a significant N550-P900 complex in the AC argues against Models 1 and 3, which both posit AC generating a local SO. The absence of N550-P900 complex in V1, M1 and S1 further argues against Model 3, which posts the generation of widespread, simultaneous SOs. The biggest change in late components occurs in the OFC, in-line with models that propose that the SO generator is primarily in orbitofrontal regions, consistent with Models 2 and 4.

On an exploratory basis, we also tested Hemisphere (Left and Right) as a fixed effect, as compared with the null model (*χ*^2^(1) = 9.66, p = .002). The results suggest that on average across Condition, elicited N550-P900 complex amplitude is greater in the left OFC than in the right OFC (Left-Right difference, M = 1.19, SE = 0.38, p = .002).

Together, the data also suggest that regional contributions to the global cortical ERP differ by brain state (Wake vs. Sleep), with earlier components (as measured via the amplitude of P200) being associated mainly with activity in the auditory cortex and to a lesser extent other regions including the OFC, and later EEG components (as measured via the amplitude of the N550-P900 complex) generated primarily in the OFC. For subsequent analyses of evoked oscillations and functional connectivity, we focused on the OFC and AC regions, as they seem most affected by brain state and are most relevant to evaluating evidence for and against the remaining SO generation models (i.e., 2 and 4; Figure 1).

#### Functional connectivity between auditory and orbitofrontal cortex following sound stimulation in sleep

The results of the connectivity analysis are represented schematically in Figure 7. LME analyses were conducted to investigate functional connectivity between AC and OFC in successive non-overlapping frequency bands (0.1-1.5 Hz, 2-4 Hz, 5-7 Hz, 8-10 Hz, 11-17 Hz, 18-29 Hz, 30-58 Hz). Functional connectivity was computed separately in each hemisphere. The mean percentage of retained epochs after outlier removal across subjects was similar in both hemispheres (Left, M = 98.7% SD = 0.9 and Right, M = 98.8%, SD = 0.6).

**Fig. 7.**
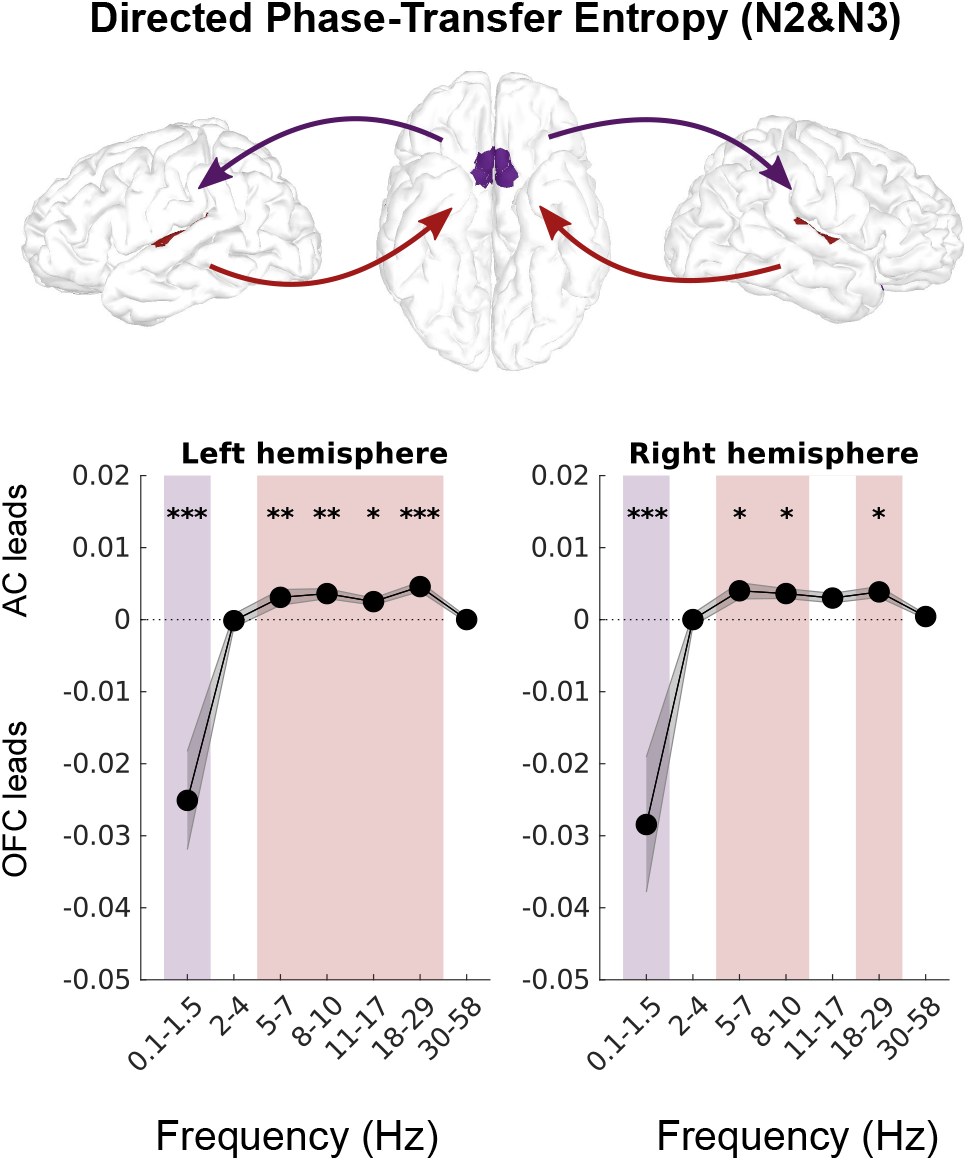
Functional connectivity (directed phase-transfer entropy) between auditory (red) and orbitofrontal (purple) cortices. Frequency bands that have significant directed connectivity are shaded in the colour of the leading region of interest. Asterisks denote significance (**\*** p < .05,**\*\*** p < .01,**\*\*\*** p < .001). N2-3: non-rapid eye movement sleep stage 2 and 3; AC: auditory cortex; OFC: orbitofrontal cortex

In both the right and left hemisphere, directed phase-transfer entropy values were significantly lower than 0 in the 0.1-1.5 Hz frequency band, indicating connectivity from OFC to AC (Left: *β* = -2.22e-02, SE = 6.89e-04, t(5.99) = -32.24, p *<* .0001; Right: *β* = -2.46e-02, SE = 9.54e-04, t(5.53) = -25.77, p *<* .0001). In the higher frequency ranges, the pattern is reversed, with modest but statistically significant directional connectivity from AC to OFC (between 5 Hz and 29 Hz on the left and similarly for the right side except that the 11-17 Hz bin was non-significant; see Figure 7 for trends and Supplementary Table 1 and 2 for numerical values).

Together these results provide evidence for an OFC to AC directed connectivity in the slow oscillation frequency band. This indicates that the N550-P900 component is generated in the OFC before propagating to other cortical areas, including the AC. These functional connectivity measurements in the low frequecies therefore are consistent with Models 2 and 4. However, we cannot rule out Model 2 at this stage, due to the possibility that information carried from AC to OFC in higher frequency bands stimulates a slow oscillation in OFC upon its arrival (which would be mediated by the lemniscal pathway). After exploring the relationship between the single-channel EEG timeseries and regional extracted time series from MEG in the next section, we will present a final analysis that is informative to distinguish Models 2 and 4.

### Effect of regionally-specific cortical states on responses

#### Investigation of region of interest contributions to EEG up-states

Closed-loop auditory stimulation usually targets EEG upstates (i.e., peaks in the slow frequency range of about 0.1-2.0 Hz) using single-channel scalp EEG, on the assumption that SO up-states in EEG are indicative of global peaks in cortical tissue excitability. In the following analyses we take advantage of MEG’s spatial and temporal resolution to explore the specificity of up-state slow wave activity as measured in single-channel scalp EEG. These analyses are intended to provide researchers using EEG for CLAS with additional information concerning the neural originals of the scalp-recorded signal, and may aid efforts to optimize stimulation effectiveness (e.g., (Navarrete et al., 2020)).

To compare the EEG recordings to the signals extracted from the AC and OFC ROIs, we first detected slow oscillation up-states in the EEG signal and defined 4-second windows (+*/ −* 2 s) around each detected peak. We then computed (zero phase lag) correlations between the EEG and each of the four source-space MEG time series (rAC, lAC, rOFC and lOFC). The average time series for EEG and the ROIs are presented in Figure 8(top). Consistent with the assumption of most CLAS studies, when the EEG is in an up-state, AC and OFC also generally seem to be in an up-state.

**Fig. 8.**
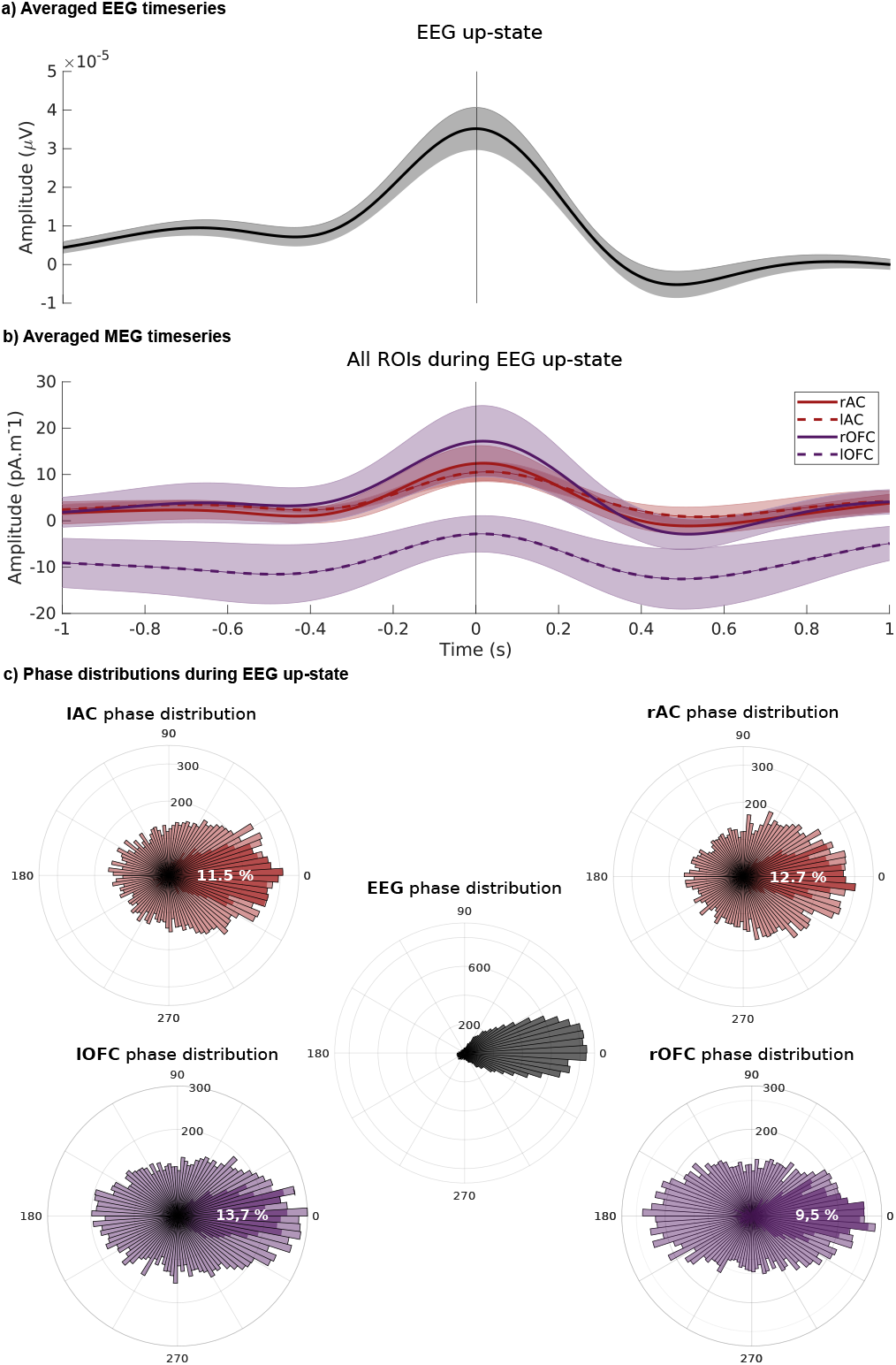
Comparison of EEG (Cz) and MEG activity from auditory (lAC, rAC) and orbitofrontal (lOFC, rOFC) regions of interest. a) EEG averages across subjects, aligned by up-state as detected in EEG. b) MEG ROI averages across subjects, aligned by up-state as detected in EEG. Coloured lines show means across participants; shaded areas indicate standard error. c) Polar histograms indicating phase distribution of EEG and ROIs during up-state as detected in EEG. The percentage of trials with aligned phase between each ROI and EEG is indicated schematically in darker shading, and numerically in superimposed white text (+*/ − <u>^π^</u> rad*). Shaded bars indicate SEM in a) and b). EEG: electroencephalography; MEG: magnetoencephalography; AC: auditory cortex; OFC: orbitofrontal cortex; r and l refers to the right and left hemispheres.

We conducted LME analyses using subjects as a random effect to identify which ROI’s activity correlates most with scalp EEG recorded at Cz, a commonly used electrode placement in SO-CLAS paradigms. Due to the intrinsic property of correlation values to be bounded between -1 and 1, no outlier removal process was implemented. Using a null-intercept linear mixed effect model with ROI as fixed effect and Subject as random effect, we found that correlations of both AC (*β* = 0.11, SE = 0.02, t(5.02) = 4.87, p = .005) and OFC (*β* = 0.08, SE = 0.02, t(5.02)= 3.35, p = .02) activity with EEG activity were significantly greater than 0 across subjects. Using a new linear mixed-effect model to compare ROI (Correlation *∼* 1 + ROI + (1 | Subject), we found a main effect of ROI. The EEG-OFC correlation was significantly lower than EEG-AC correlation (*β* = -3.49e-02, SE = 2.68e-03, t(6.590e+04) = -13.03, p < .0001). The strong representation of AC in this EEG montage is consistent with the scalp distribution of brain activity generated in auditory cortex (Stropahl et al., 2018); frontal activity may be less well represented due to distance from sensors or tissue orientation (Ahlfors et al., 2010).

We also further explored the lateralization effect noted in the previous analyses. Adding Hemisphere as a second fixed effect increased model fit (*χ*^2^(1) = 483.62, p < .0001), with overall EEG activity being more correlated with ROI activity in the left hemisphere (F(1, 65899) = 485.40, p < .0001). This result is coherent with the previously observed dominance of the left hemisphere in other analysis.

#### Investigation of phase relationships between regions of interest and EEG up-state

To further explore the relationship between single-channel EEG and regional brain activity, we calculated the instantaneous phase of the extracted ROI signals at the time EEG up-states were detected. Because phase angles are bounded between *−π* and *π* rad, no outlier removal process was implemented. As shown in Figure 8 (bottom), phase distribution from the four key regions of interest were far more dispersed than in the EEG, and only a small percentage of trials were aligned with a simultaneous EEG up-state (phase difference between EEG and ROI +*/ − <u>^π^</u> rad*; lAC M = 11.5 % SD = 0.9; rAC M = 12.7 %, SD = 1.6; lOFC M = 13.7 %, SD = 4.6; rOFC M = 9.5 %, SD = 1.7). We then computed F-scores to compare the variance of the EEG distribution with that of each ROI. The variance of phase distribution (i.e., dispersion) of each ROI was significantly greater than that of EEG (F(16487) = 0.33, p < .001), lAC phase distribution (F(16487) = 0.33, p < .0001), lOFC phase distribution (F(16487) = 0.26, p < .001) and rOFC phase distribution (F(16487) = 0.31, p < .001). These results indicate that up-states in EEG only partially coincide with local up-states in key cortical regions.

#### Comparison of EEG evoked responses as a function of regional tissue state

Returning to the main research questions concerning models of information flow leading to evoked slow oscillations, we now investigate the effect of tissue excitability on amplitude of the evoked response, a measure of stimulation success. Two pieces of information from previous analyses make this analysis relevant: first, evoked N550-P900 amplitude is normally distributed (see Supplementary Figure 2), indicating that auditory stimulation varies on a trial-to-trial basis in its effectiveness; this observation suggests room for optimization. Second, because up-states in EEG only partially coincide with local up-states in key cortical regions (Figure 8), it is possible that stimulation effectiveness could depend on the phase of tissue in specific regions.

To identify the impact of regionally-specific cortical tissue excitability level on the evoked brain response, we take advantage of the open-loop nature of the study design, which allows us to observe the effects of stimulations occurring over the full cycle of tissue excitability. We detected up-states and down-states of slow activity (0.1 - 2.0 Hz) in each key region of interest. We then computed the amplitude of the elicited N550-P900 complex and spindle activity across condition (Up-state vs. Downstate). We focused on evoked responses as measured in EEG for comparability with previous literature. Concerning the elicited N550-P900 complex amplitude, we conducted LME analyses to investigate the impact of up and down-states of both the OFC and the AC.

Including Condition (Up-state vs. Down-state) as a fixed effect in a linear mixed-effect model increased model fit (Amplitude *∼* 1 + Condition +(1|Subject)) as compared to a null intercept-only model (*χ*^2^(1) = 4.08, p = .043), indicating across ROIs, stimulation occurring during an up-state elicited a larger-amplitude N550-P900 complex as observed in EEG, compared to down-states. Adding the interaction of ROI and Condition (Amplitude *∼* 1 + Condition*ROI +(1|Subject)) resulted in a better-fitting model (*χ*^2^(2) = 15.34, p = .0005). Post-hoc analysis using estimated means showed that tissue state in the OFC was critical to the generation of N550-P900 complex (OFC: Up vs. Down-state difference, M = 8.43, SE = 2.01, p = .0002), whereas tissues state in the AC did not significantly affect N550-P900 complex amplitude as measured in EEG (AC: Up vs. Down-state difference, M = -2.56, SE = 1.96, p = 0.56).

These results provide additional evidence for a critical role of the OFC in SO generation. Auditory cortex excitability state instead does not seem to matter, further implicating the non-lemniscal auditory pathway in the CLAS effect and decreasing the likelihood of Model 2, in which the auditory information passes first to the AC via the lemniscal pathway. Together, these results are best explained by Model 4.

On an exploratory basis, we also tested Hemisphere (Left and Right) as a fixed effect, as compared with the previous model (*χ*^2^(1) = 0.03, p = 0.86). The comparison suggests that Hemisphere does not account for more variance in the data, meaning that differences in tissue state across left and right OFC does not clearly impact the N550-P900 complex as observed in EEG. Recalling the importance of elicited sleep spindles in the memory consolidation effects of CLAS (Fernandez and Lüthi, 2020, Harrington and Cairney, 2021, Ngo et al., 2013), and that stronger N550-P900 complexes are related to greater evoked spindle activity (Figure 5b), we investigated the effect of cortical state in ROIs on spindle activity. For each epoch, root mean squared values of the EEG sigma filtered in the previously defined frequency band of interest (11-14 Hz) was computed in the time windows of interest (0.6 to 1.2 s post-stimulation), as well as before stimulation to serve as baseline (-0.6 to 0 s). Note that these analyses do not assist us in distinguishing between models of information flow leading to generation of evoked slow oscillations, but as spindles are integral to memory consolidation processes, they are nonetheless helpful to better understand the CLAS effect.

We first investigated the conditions affecting the production of spindle power via auditory stimulation, using zero-intercept models (Spindle Power Change *∼* 0 + ROI + (1 | Subject) and Spindle Power Change *∼* 0 + Condition + (1 | Subject)). In each ROI, across Condition (Up and Down-state merged), auditory stimulation statistically increased spindle power as compared to a pre-stimulation baseline (AC, *β* = 0.51, SE = 0.19, t(5.3522) = 2.74, p = .04 and OFC, *β* = 0.55, SE = 0.19, t(5.42) = 2.82, p = .04). Collapsing instead across ROIs, auditory stimulation statistically increased spindle power in Up-states (*β* = 0.58, SE = 0.18, t(5.31) = 3.16, p = .02), but not Down-states (*β* = 0.44, SE = 0.19, t(5.51) = 2.35, p = .06).

To compare between ROIs, we used a LME with Subjects as random effect (Spindle Power Change *∼* 1 + Condition + (1 | Subject)). We find a main effect of Condition (tissue excitability as measured with ROI Up and Down-state, (F(5716) = 4.14, p = .04)) with Up-state stimulation eliciting greater spindle power, as measured in EEG, than Down-state stimulation. Adding the interaction between ROI and Condition (Amplitude *∼* 1 + Condition*ROI +(1|Subject)) did not improve model fit (*χ*^2^(2) = 2.18, p = 0.34). These results show that while being in an up-states in either regions of interest when sound arrives is associated with greater evoked spindle activity, we do not observe regional dependencies.

For completeness, we also tested Hemisphere (Left and Right) as a fixed effect (Spindle Power Change *∼* 1 + Condition + Hemisphere + (1 | Subject)), as compared with the Condition-only model (*χ*^2^(1) = 1.69, p = 0.19). The comparison suggests that Hemisphere does not account for additional variance in the data, meaning that differences in tissue state across left and right OFC do not clearly impact spindle generation.

## Discussion

### Evaluating evidence for SO generation models

Understanding the neurophysiology of closed-loop auditory stimulation requires identification of the origin of auditory evoked brain responses during sleep. The main goal of this work was to define potential mechanistic models based on previous literature (Figure 1) and to evaluate their ability to explain spatially-resolved evoked brain responses in the sleeping brain, including slow oscillations and sleep spindles, and by association, the beneficial effects on memory consolidation that have been reported in literature (Choi et al., 2020, Harrington and Cairney, 2021, Nasr et al., 2022), as well as other sleep-related physiological functions (Besedovsky et al., 2017, Kelley et al., 2022).

According to the first potential model we described (Figure 1), slow oscillations would be generated locally within the auditory cortex upon the arrival of sound information through lemniscal auditory pathways at primary auditory cortex in the superior temporal gyrus. If this were the case, we would expect SOs to occur first in this region before propagating to subsequent cortical regions. Although we do observe clear evoked auditory responses in sleep that are similar to those in the Wake state up to and including 200 ms (P200), the evoked slow oscillations appear strongest in the orbitofrontal region (Figure 4, 6). Additional evidence against this model comes from the functional connectivity analysis (see Figure 7), which shows that in the slow oscillation frequency range (0.5-1.5 Hz), information flows in the reverse direction; from OFC to AC (replicated in both left and right hemispheres). These findings align with previous reports that the primary generator of (endogenous) SOs is found in ventral frontal areas (Morgan et al., 2021).

In the second of the proposed models (Model 2 in Figure 1), sound information would arrive at the auditory cortex via primary auditory pathways and would then be passed forward to frontal regions, which would generate slow oscillations. This model relies on the progression of auditory information from auditory cortex to ventral frontal regions. Although we did observe connectivity from AC to OFC in higher frequencies (i.e., *>* 5Hz), it was of very low strength. The majority of the energy in auditory evoked responses is captured in the *<* 5 Hz frequency range (confirmed by applying filters with different band pass limits). Although it cannot entirely be ruled out on the basis of the present analysis, it seems unlikely that the modest AC to OFC connectivity we observed in high frequency ranges is the trigger for SO generation in OFC. Additionally, the finding that early auditory evoked components in AC and OFC are differentially impacted by sleep state, with amplitude in AC decreasing and OFC increasing, suggests that auditory information travelling through lemniscal pathways to AC may not be the most critical to activate SOs within OFC. Moreover, the observation that SOs are more likely to be generated when tissue in OFC is in an up-state at the time of stimulation as compared with AC, suggests that the non-lemniscal auditory pathway is most relevant for the CLAS effect.

Models 3 and 4 consider the involvement of the non-lemniscal pathway as a potential mechanism for auditory evoked responses as suggested by Bellesi et al. (2014). They hypothesized that CLAS is likely mediated by the activation of the non-lemniscal ascending auditory pathway, which projects broadly to association areas including frontal regions and secondary auditory areas (Bellesi et al., 2014, Kjaerby et al., 2022). They proposed that the modulation of noradrenaline levels by the locus coeruleus is critical for the generation of N550-P900 complexes, and that low noradrenaline levels might be a necessary factor for the emergence of an evoked N550-P900 complex, whereas elevated levels of noradrenaline would result in arousal.

In Model 3, a signal generated in the arousal network via interactions with the non-lemniscal auditory system in the brainstem and thalamus would cause changes to neural activity in widespread areas, and generate slow oscillations throughout the cortex. Although this domain-generality is plausible from work showing that visual, tactile, and auditory stimulation can all generate slow oscillations (Riedner et al., 2011), we do not find that slow oscillations are generated via auditory stimulation in any of the other selected cortical regions (V1, M1, S1), nor do we observe activity within the whole brain maps during the evoked N550-P900 complex (Figures 4 and 6). Moreover, the significant functional connectivity in the slow wave frequency band from OFC to AC would not be observed if both the OFC and the AC were generators of slow oscillation receiving simultaneous stimulation from a common source (Figure 7).

Finally, in Model 4, the general arousal system is implicated, but it is the ventral frontal regions that generate slow oscillations. This model is supported by the finding that frontal and ventral regions show the strongest SOs (Figures 4, 3, 6), as well as the functional connectivity analysis, which shows that information flows from OFC to AC bilaterally in the slow wave frequency range (Figures 7). Furthermore, the result that tissue excitability state in OFC significantly affects the success of stimulation to generate a strong N550-P900 complex but AC does not, further supports Model 4 (Figures 9).

**Fig. 9.**
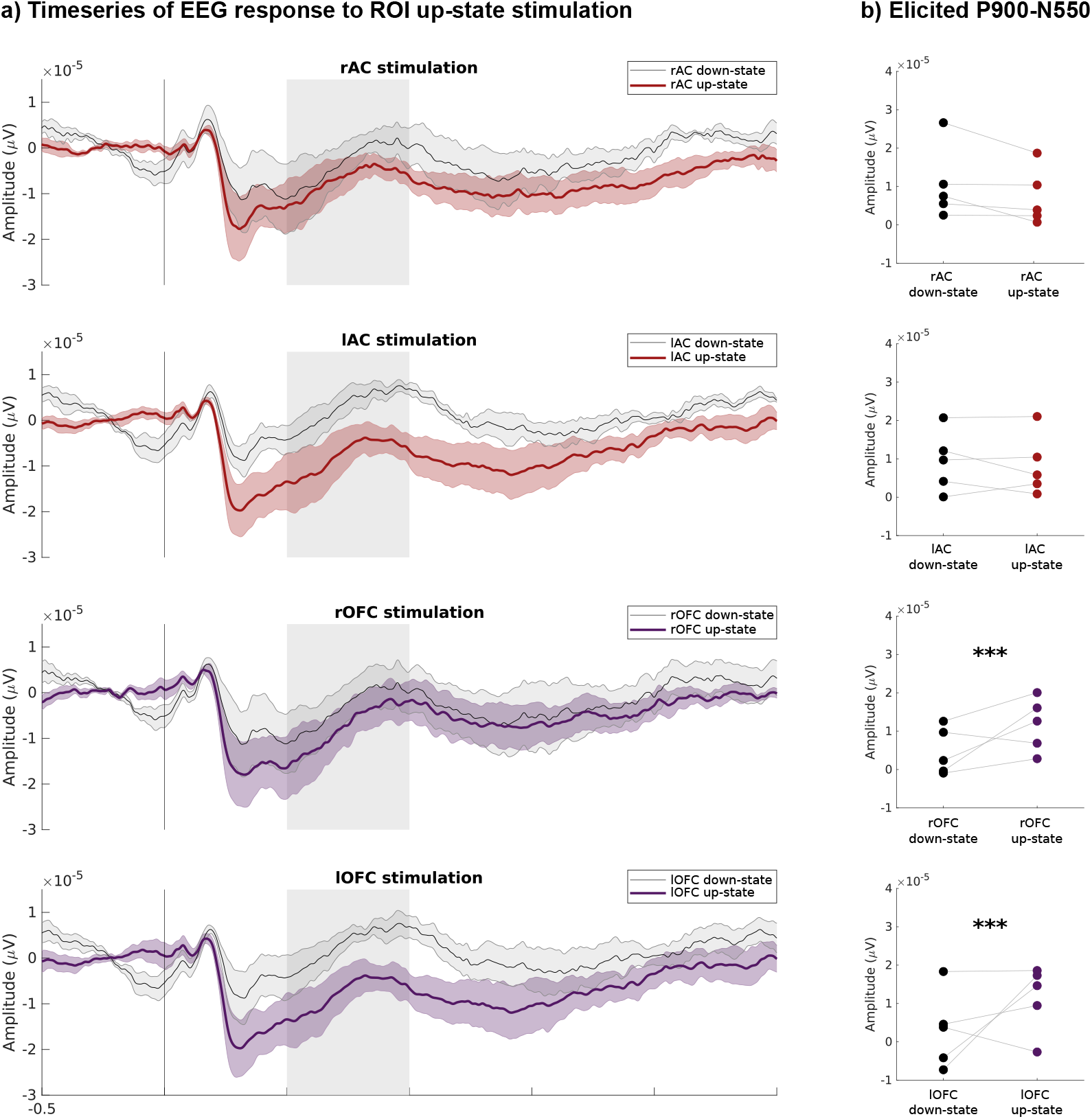
Evoked responses in EEG divided according to the excitability state of local tissue in auditory (rAC, lAC), and orbitofrontal (rOFC and lOFC) regions of interest, as measured in MEG. a) Comparison of the EEG time series for up-state and down-state stimulations, which have been defined by the state of excitability in MEG extracted timeseries from auditory and orbitofrontal ROIs. Coloured lines show means across participants; shaded areas indicate standard error. Grey shading indicates the evoked responses components P200, N350 and N550-P900. b) Mean N550-P900 complex amplitude for each ROI in each condition. Each subject is represented by its mean. Asterisks denote significant differences between up-state and down-state stimulation using LME statistical models (*** p < .001). MEG: magnetoencephalography; AC: auditory cortex; OFC: orbitofrontal cortex; r and l refers to the right and left hemispheres.

### Evoking sleep spindles

Although the primary objective of this work is to evaluate evidence in support of the four models of information flow leading to the generation of evoked slow oscillations, we also investigated the generation of spindles evoked by auditory stimulation, due to their importance in sleep’s memory consolidation function and in the CLAS effect (Fernandez and Lüthi, 2020, Ngo et al., 2013). Previous research focused on the increase in spindle activity when auditory stimulation is time-locked to the up-state of the targeted slow oscillation (Ngo et al., 2013). However, a study by (Sato et al., 2007) showed that non-arousing stimulation using different sensory modalities (somatosensory, auditory, or visual) during light non-rapid eye movement (stage N2) sleep also induced higher spindle activity in related sensory cortical areas. Spindle generation is often not reported in sleep studies focusing on ERPs (Colrain and Campbell, 2007), as they are not precisely time-locked to stimulation events and disappear when independent trials are averaged (in the time domain). By using time-frequency analyses and analyzing sigma power within specific time windows following stimulation, we replicated the observation that sleep spindles are elicited in N2 and N3 sleep (Ngo et al., 2013). Furthermore, we showed that spindle amplitude was bigger in trials that successfully generated higher N550-P900 complexes. MEG topographies of these evoked spindles show a pattern of activation in medial and dorsal brain regions including the precuneus, which is similar to the pattern of activity in the spindle frequency range throughout N2 and N3 sleep stages (see Figure 5).

Interestingly, even if their presence is linked to the amplitude of the evoked N550-P900 complex (as seen the Figure 5b), the elicited spindle activity does not seem to depend on the local activity of the key regions of interest we selected for our analysis; both AC and OFC up-state stimulation induces an increase in spindle activity which is greater than down-state stimulation. We can hypothesize that even if the spindle and the N550-P900 generation mechanisms are intertwined, they are not identical. Other factors, potentially the state of activity in other regions, might modulate the degree of spindle activity generated. This observation opens the possibility that the two mechanisms might be independently manipulated, which could be useful for restoring ideal functional coupling between SOs and sleep spindles, which degrades with age and is linked to memory consolidation (Hahn et al., 2020, Helfrich et al., 2018).

### Improving the effectiveness of CLAS

While the effect of CLAS on physiology and memory largely replicates across studies (Choi et al., 2020, Harrington and Cairney, 2021, Wunderlin et al., 2021), there are inter-individual differences in the effectiveness of non-invasive brain stimulation (Guerra et al., 2020, Ziemann and Siebner, 2015), and effectiveness appears to change over the lifespan (Schneider et al., 2020). Attempts have therefore been made to identify the best timing for CLAS with respect to tissue excitability state as detected in a single EEG channel (in younger and older populations) (Navarrete et al., 2020), and to target local SOs in different brain regions using the spatial information that is available in multichannel EEG data (Ruch et al., 2022). Our investigation of phase in EEG and MEG makes several contributions to this area. Notably, we found that although EEG up-states detected in single channel over frontal areas does on average correlate with up-states in both AC and OFC (Figure 8a), only *∼*10-15% of slow oscillation up-states detected in EEG correspond to up-states in each of these regions (Figure 8b). As the state of the OFC is most critical to successful generation of N550-P900 complexes and their associated spindles (Figure 9), which both play important roles in memory consolidation, particularly when they are coupled (Hahn et al., 2020, Helfrich et al., 2018, Mikutta et al., 2019, Muehlroth et al., 2019), it is likely that increased efficiency and reliability of CLAS can be obtained by selectively targeting up-states in ventral frontal regions. This goal could be achieved by detecting patterns in multichannel EEG (e.g., (Ruch et al., 2022)), and/or machine learning-approaches that can detect fine-grained signatures on one or several channels to stimulate the brain with optimal timing (e.g., Valenchon et al. (2022)), and perhaps adapt detection parameters in real time to an individual’s idiosyncratic neural signals. These ideas are in alignment with the emerging concepts of ‘precision neuroimaging’ and ‘person specific methods’, in which reliable individual differences in brain activation or connectivity are studied to understand (and sometimes make use of) individual differences in brain function and behaviour Michon et al. (2022).

As regards the origins of the auditory evoked responses in sleep, our results suggest the following contributions: the first components (i.e., P100, P200) originate mainly in bilateral auditory cortex (as expected (Stropahl et al., 2018)), perhaps with some contribution from other brain regions including OFC via non-lemniscal pathways. Later components of the observed brain response in the EEG appear to come mainly from the OFC upon its activation through a non-lemniscal pathway (Figure 4).

### Limitations and next steps

A limitation of our investigation of is that while we have evidence against Models 1 and 3 and there is generally stronger evidence for Model 4 as best explaining information flow leading to the generation of evoked slow oscillations, we cannot entirely rule out Model 2, because we do observe some feed-forward activity from AC to OFC in the higher frequency ranges (i.e., above 5 Hz). Notably, auditory information does not pass through the lemniscal *or* non-leminiscal branches, but rather both on its way to other cortical regions, in both sleep and wakefulness. It seems likely that the small amount of connectivity we observed from AC to OFC may be capturing information flow that is unrelated to SO generation in OFC. However, invasive methods would be required to confirm this hypothesis. For example, if auditory stimulation continued to generate SOs in OFC when bilateral AC had been ablated or disconnected would provide definitive evidence against Model 2. Another limitation of our study is that due to the nature of its design, we were not able to distinguish between elicited (evoked) and endogenous brain activity. The topographies of endogenous SOs and spindles offered in Figures 3b and 5d for visual comparison with the evoked activity are therefore approximations that are strongly weighted towards endogenous activity, but nonetheless also capture evoked activity. Comparing periods of evoked oscillations in stimulated sleep and endogenous oscillations in unstimulated sleep in the same individuals would allow for a more rigorous evaluation of the similarities and differences in slow oscillations and spindles generated by sound (which was not an aim of the present work). The current work does not empirically address how generating slow oscillations through sound stimulation causes the memory consolidation processes that have been observed in previous work Choi et al. (2020), Harrington and Cairney (2021), Ngo et al. (2013, 2015), because we did not have a memory-related experimental manipulation. The apparently common origin of evoked and endogenous slow oscillations in ventral frontal brain regions Morgan et al. (2021), Riedner et al. (2011) suggests that memory consolidation processes are activated through stimulation in the same way as they occur spontaneously (see Brodt et al. (2023), Staresina et al. (2023)). However, direct comparison of spontaneous vs. evoked memory processes is needed to confirm this assumption.

Nonetheless, the study aims to investigate the evoked responses which have been causally implicated in memory processes in previous work (e.g., (Harrington and Cairney, 2021, Ngo et al., 2013)). The overnight design allowed us to record data in all sleep stages and have large numbers of trials in N2 and N3 sleep, which is difficult to achieve in a nap design. Our LME-based statistical approach allowed us to investigate our research questions within a small sample size, but as we only record data in five participants, we cannot account for the range of human variability in brain responses to sound in sleep (limiting generalizability). It would be interesting to explore individual differences across populations and over the lifespan, in a larger sample. This research direction could help to explain why sensitivity to CLAS differs between individuals (Guerra et al., 2020, Ziemann and Siebner, 2015) and in older adults (Navarrete et al., 2020).

Several of our analyses suggest a leftward lateralization in OFC involvement. We caution against over-interpreting these findings, as hemispheric differences in brain folding and ROI placement can affect the strength of observed signals in MEG source space; however, other studies using EEG have also found that low frequency oscillations in ventral limbic areas are more frequent on the left side (Achermann et al., 2001, Morgan et al., 2021), suggesting that a more focused analysis of lateralization may be an interesting focus for future work. Finally, we opted to use standard surface-based cortical source localization models. Previous work has demonstrated that brain signals associated with information in the auditory lemniscal pathway can be extracted using MEG source space volume models (Coffey et al., 2016); however, signals extracted from these models are more difficult to work with and their use with functional connectivity metrics has not been validated. As MEG modelling and connectivity techniques develop, it may be possible to explore the involvement of brainstem regions and more directly test the involvement of the ARAS in brainstem-OFC interactions, to further refine models of SO generation. As we collectively push the limits of methodology, preregistration may facilitate reproducible results and reduce publication bias (Logg and Dorison, 2021).

## Conclusion

In summary, our data suggest a crucial orbitofrontal contribution to the auditory-evoked N550-P900 complex that underlies the effect of closed-loop auditory stimulation on physiology and memory. Previous research mainly detected high states of excitability based on single channel scalp EEG, making the assumption that activity over frontal regions represents a global up-state of high brain tissue excitability. Our results suggest that targeting up-states in ventral frontal regions has potential to improve the effectiveness of CLAS. Clarifying how sound stimulates the brain has important implications for making causal manipulation tools like CLAS more effective in fundamental research, and in potential clinical applications.

### Data availability statement

The raw MEG and EEG sleep recordings are accessible via the OMEGA open MEG archive (Niso et al., 2016). All extracted values used for statistical analyses are available on the Open Science Framework website (https://osf.io/afv9g/?view_only=24576c983b8941b1b429e18b42e17061).

## Author contributions

CRediT roles: Hugo Jourde: Methodology, Formal analysis, Writing - Original Draft, Writing - Review & Editing, Visualization. Raphaëlle Merlo and Meredith Rowe: Data collection, Writing - Review & Editing. Mary Brooks: Writing – Original Draft, Writing - Review & Editing. Emily Coffey: Conceptualization, Methodology, Formal analysis, Data Curation, Writing - Original Draft, Writing - Review & Editing, Visualization, Supervision, Project administration, Funding acquisition.

## Acknowledgements

The authors would like to thank Sylvain Baillet and Mark Lalancette for assistance with MEG technical matters, Robert Zatorre for assistance with funding acquisition and consultation concerning results, Alexander Albury, Charlotte Emma Moore, Keelin Greenlaw and Marie-Anick Savard for help with statistical modelling, Marie-Anick Savard and Anita Paas for helpful comments on the manuscript, Alix Noly-Gandon for helping with several data recordings and confirming the sleep scoring, and lab members for general support. This work was financially supported by a Natural Sciences and Engineering Research Council of Canada (NSERC) Discovery grant, a grant from the Fonds de recherche du Québec – Nature et technologies (FRQNT), the ‘Healthy Brains for Healthy Lives’ Canada First Research Excellence Fund (CFREF) at McGill University, and a Concordia (New Scholar) Research Chair in ‘Sleep and Sound’ to EBJC. HRJ was supported by a Graduate Scholar Stipend from the Centre for Research on Brain, Language and Music.

