## Supplementary Materials for "The neurophysiology of closed-loop auditory stimulation in sleep: a magnetoencephalography study"

[1]Concordia University, Montreal, Canada
[2]International Laboratory for Brain, Music and Sound Research (BRAMS)
[3]Centre for Research on Brain, Language and Music (CRBLM)
[4]Quebec Bio-Imaging Network (QBIN)
[5]Université de Montréal, Montreal, Canada
[6]McGill University, Montreal, Canada

### Timeseries of auditory evoked responses (MEG)

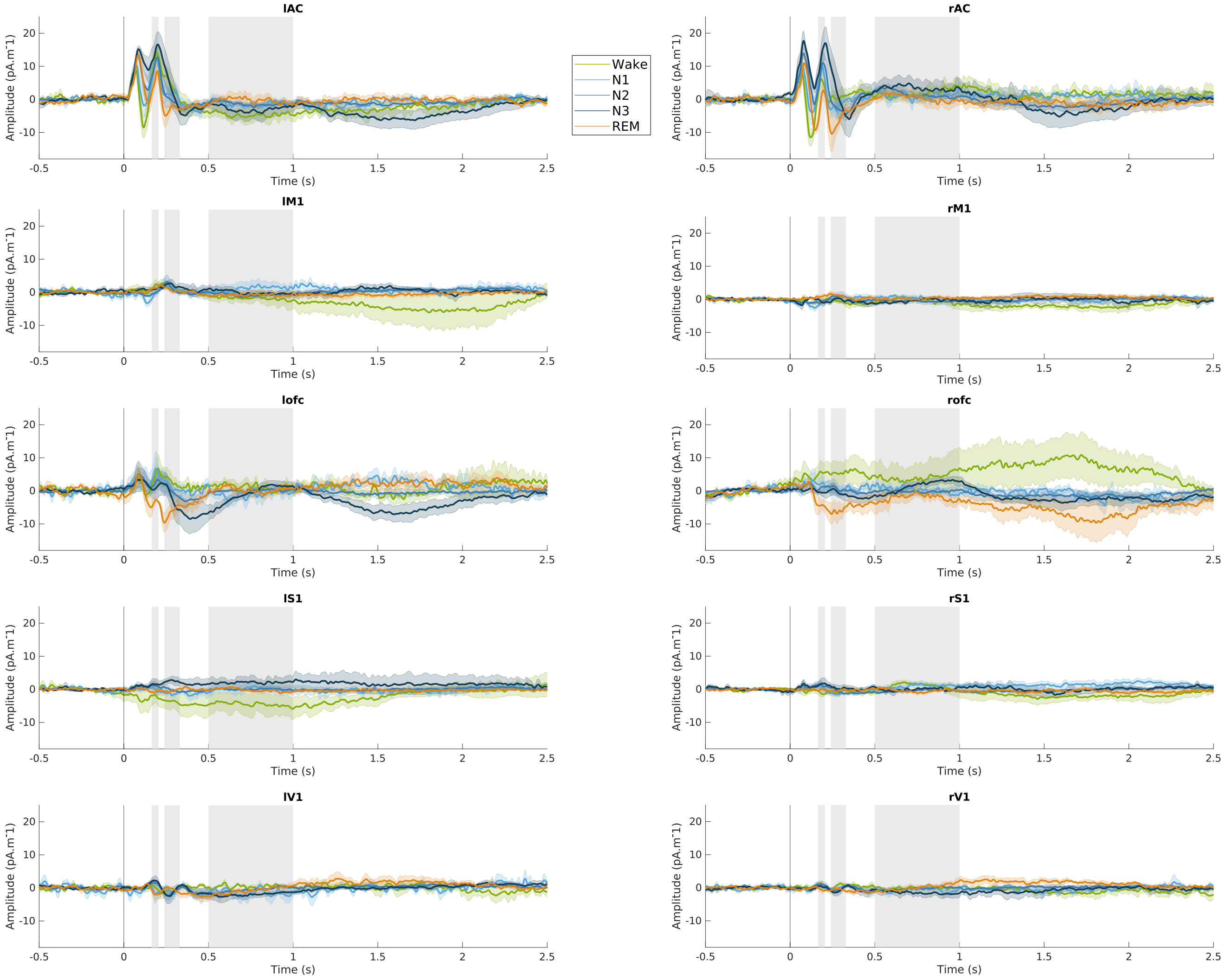


**Fig. 1.** Evoked responses across sleep stages. Time series, averaged across subjects, for each region of interest. Coloured lines show means across participants; shaded areas indicate standard error. Grey shading indicates the evoked responses components P200, N350 and N550-P900.

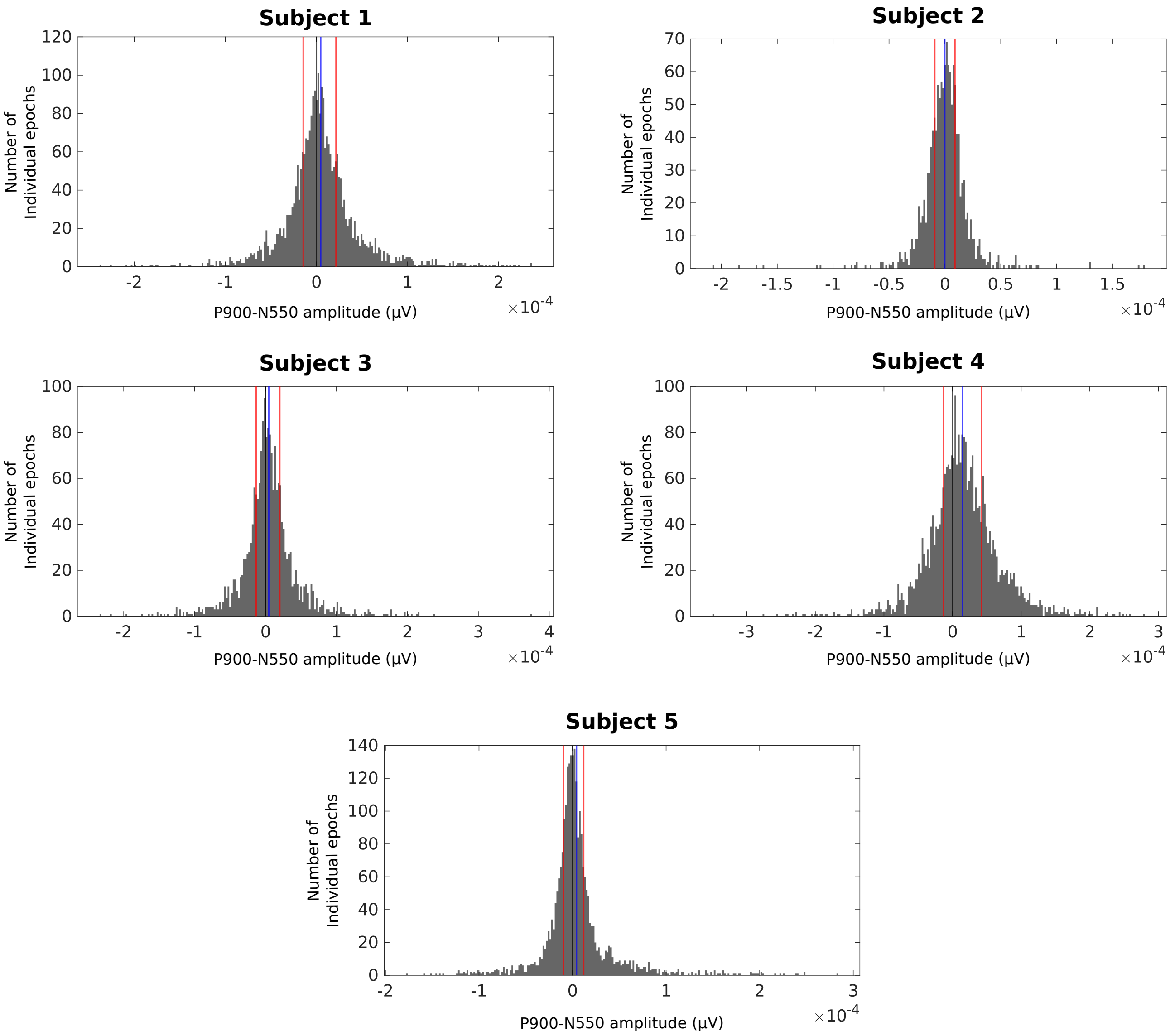


**Fig. 2.** Histograms of P900-N550 amplitude for each subjects. Black lines represent 0, blue lines represent the average value for each subject. Red lines represent the Top vs. Bottom 25% for each subject

**Table 1.** Fixed effects of the connectivity analysis for the left hemisphere. Negative estimate values indicate OFC leading and positive estimate values indicate AC leading. Degrees of freedom (df) are approximate and estimated using Satterthwaite's method.

| Fixed effects: | Estimate | Std. Error | df | t value | Pr(>ltl) |
|---|---|---|---|---|---|
| FrequencyBand1 (0.1-1.5 Hz) | -2.22e-02 | 6.89e-04 | 5.98 | -32.236 | 6.06e-08 |
| FrequencyBand2 (2-4 Hz) | -5.547e-04 | 6.89e-04 | 5.96 | -0.805 | 0.45150 |
| FrequencyBand3 (5-7 Hz) | 2.83e-03 | 6.89e-04 | 5.96 | 4.106 | 0.00640 |
| FrequencyBand4 (8-10 Hz) | 3.28e-03 | 6.89e-04 | 5.96 | 4.759 | 0.00318 |
| FrequencyBand5 (11-17 Hz) | 2.21e-03 | 6.89e-04 | 5.96 | 3.214 | 0.01843 |
| FrequencyBand6 (18-29 Hz) | 4.15e-03 | 6.89e-04 | 5.96 | 6.033 | 0.00096 |
| FrequencyBand7 (30-58 Hz) | -3.31e-04 | 6.89e-04 | 5.96 | -0.481 | 0.64784 |

**Table 2.** Fixed effects of the connectivity analysis for the right hemisphere. Negative estimate values indicate OFC leading and positive estimate values indicate AC leading. Degrees of freedom (df) are approximate and estimated using Satterthwaite's method.

| Fixed effects: | Estimate | Std. Error | df | t value | Pr(>\|t\|) |
|---|---|---|---|---|---|
| FrequencyBand1 (0.1-1.5 Hz) | -2.46e-02 | 9.54e-04 | 5.53 | -25.78 | 5.62e-07 |
| FrequencyBand2 (2-4 Hz) | -3.45e-04 | 9.54e-04 | 5.51 | -0.36 | 0.7307 |
| FrequencyBand3 (5-7 Hz) | 3.50e-03 | 9.54e-04 | 5.51 | 3.68 | 0.0121 |
| FrequencyBand4 (8-10 Hz) | 3.11e-03 | 9.54e-04 | 5.51 | 3.26 | 0.0195 |
| FrequencyBand5 (11-17 Hz) | 2.45e-03 | 9.54e-04 | 5.51 | 2.57 | 0.0456 |
| FrequencyBand6 (18-29 Hz) | 3.31e-03 | 9.54e-04 | 5.51 | 3.48 | 0.0151 |
| FrequencyBand7 (30-58 Hz) | -5.02e-05 | 9.54e-04 | 5.51 | -0.05 | 0.9599 |
